# TARANIS interacts with VRILLE and PDP1 to modulate the circadian transcriptional feedback mechanism in *Drosophila*

**DOI:** 10.1101/2023.05.19.541420

**Authors:** Oghenerukevwe Akpoghiran, Dinis J.S. Afonso, Yanan Zhang, Kyunghee Koh

## Abstract

The molecular clock that generates daily rhythms of behavior and physiology consists of interlocked transcription-translation feedback loops. In *Drosophila*, the primary feedback loop involving the CLOCK-CYCLE transcriptional activators and the PERIOD-TIMELESS transcriptional repressors is interlocked with a secondary loop involving VRILLE (VRI) and PAR DOMAIN PROTEIN 1 (PDP1), a repressor and activator of *Clock* transcription, respectively. Whereas extensive studies have found numerous transcriptional, translational, and post-translational modulators of the primary loop, relatively little is known about the secondary loop. In this study, using male and female flies as well as cultured cells, we demonstrate that TARANIS (TARA), a *Drosophila* homolog of the TRIP-Br/SERTAD family of transcriptional coregulators, functions with VRI and PDP1 to modulate the circadian period and rhythm strength. Knocking down *tara* reduces rhythm amplitude and can shorten the period length, while overexpressing TARA lengthens the circadian period. Additionally, *tara* mutants exhibit reduced rhythmicity and lower expression of the PDF neuropeptide. We find that TARA can form a physical complex with VRI and PDP1, enhancing their repressor and activator functions, respectively. The conserved SERTA domain of TARA is required to regulate the transcriptional activity of VRI and PDP1, and its deletion leads to reduced locomotor rhythmicity. Consistent with TARA’s role in enhancing VRI and PDP1 activity, overexpressing *tara* has a similar effect on the circadian period and rhythm strength as simultaneously overexpressing *vri* and *Pdp1*. Together, our results suggest that TARA modulates circadian behavior by enhancing the transcriptional activity of VRI and PDP1.

**Statement of Significance:** Internal molecular clocks generating circadian rhythms of around 24 hours broadly impact behavior and physiology, and circadian dysfunction is associated with various neurological and metabolic diseases. The *Drosophila* circadian clock is a valuable model for understanding the molecular mechanisms underlying daily rhythms as many components of the clock are highly conserved. In this study, we identify a conserved gene, *taranis*, as a novel regulator of the *Drosophila* molecular clock. We show that TARANIS modulates circadian behavior by physically interacting with and enhancing the transcriptional activity of clock proteins VRILLE and PDP1. Since mammalian homologs of VRILLE and PDP1 also function in the molecular clock, our results have implications for understanding the mammalian circadian clock.

## Introduction

The molecular mechanisms of endogenous circadian oscillators are conserved across organisms, making *Drosophila melanogaster* a valuable model for understanding the genetics of the circadian clock (Allada and Chung, 2010; Hardin, 2011; Tataroglu and Emery, 2015; Patke et al., 2020). The molecular clocks in flies and mammals consist of interlocked transcription-translation feedback loops that drive rhythmic gene expression. In the *Drosophila* clock, the CLOCK-CYCLE (CLK-CYC) heterodimer binds to the E-box elements in the *period* (*per*) and *timeless* (*tim*) promoters to activate their transcription, and the PER-TIM heterodimer inhibits CLK-CYC, suppressing their own transcription (Darlington et al., 1998). In an interlocked loop, CLK-CYC drives the transcription of *vri* and *Pdp1*. VRI and PDP1 competitively bind to the D-box elements in the *Clk* promoter to repress and activate *Clk* transcription, respectively (Cyran et al., 2003; Glossop et al., 2003). The mammalian clock involves analogous, although more complex, interlocked feedback loops, including nuclear receptors and mammalian homologs of VRI and PDP1 (Partch et al., 2014; Takahashi, 2017; Patke et al., 2020). While many molecules underlying the transcriptional, translational, and post-translational modulation of the primary feedback loop have been identified (Allada and Chung, 2010; Zheng and Sehgal, 2012; Dubowy and Sehgal, 2017), the mechanisms regulating the secondary interlocked loop are less well understood (Blau and Young, 1999; Cyran et al., 2003; Glossop et al., 2003; Benito et al., 2007; Zheng et al., 2009; Ling et al., 2012; Gunawardhana and Hardin, 2017; Gunawardhana et al., 2021).

VRI and PDP1 are bZIP transcriptional factors essential for proper development and circadian behavior (George and Terracol, 1997; Blau and Young, 1999; Reddy et al., 2000; McDonald and Rosbash, 2001; Cyran et al., 2003; Glossop et al., 2003). Null mutants of *vri* and *Pdp1* are lethal, but knocking down either gene in clock neurons leads to arrhythmic behavior despite a functioning molecular clock, suggesting their role in regulating locomotor rhythms downstream of the molecular clock (Blau and Young, 1999; Cyran et al., 2003; Gunawardhana and Hardin, 2017). Additionally, *vri* overexpression substantially lengthens the circadian period (Blau and Young, 1999), while an isoform-specific *vri* deletion mutant exhibits a shorter period (Gunawardhana et al., 2021), indicating VRI’s role in regulating the clock speed. An isoform-specific, arrhythmic *Pdp1ε* mutant exhibits reduced CLK expression (Zheng et al., 2009), demonstrating that PDP1 also functions in the molecular clock. Although PDP1 overexpression has little effect on the period length (Zheng et al., 2009), heterozygous *Pdp1* null mutants show slightly longer periods (Cyran et al., 2003), suggesting that PDP1 also modulates the clock speed.

We previously reported that TARA interacts with Cyclin A (CycA) and Cyclin-dependent kinase 1 (Cdk1) to promote sleep in *Drosophila* (Afonso et al., 2015b). In addition, TARA and its mammalian homologs, Transcriptional Regulator Interacting with PHD-Bromodomain (TRIP-Br, also known as SERTAD) family of proteins, function in diverse biological processes, including cell cycle progression, metabolic regulation, and transcriptional coregulation (Hsu et al., 2001; Calgaro et al., 2002; Liew et al., 2013; Dutta and Li, 2017). TARA was initially categorized as a Trithorax group (TrxG) member of transcriptional coactivators (Calgaro et al., 2002). However, a later study found that TARA enhances Polycomb (Pc)-mediated transcriptional repression (Dutta and Li, 2017), suggesting that TARA may act as a transcriptional coactivator or corepressor depending on the cellular context and interaction partners. Here, we report a novel role of TARA in the transcriptional coregulation of VRI and PDP1 in the molecular clock. Our results suggest that TARA modulates the pace of the molecular clock and locomotor activity rhythms by physically interacting with and enhancing the transcriptional activity of VRI and PDP1. Our study has implications for understanding the mammalian circadian clock since mammalian homologs of VRI and PDP1 also function in the molecular clock.

## Materials and Methods

### Fly stocks

Flies were raised on standard food containing molasses, cornmeal, and yeast at 25°C under 12 h-12 h light-dark (LD) cycle. The following lines were obtained from the Bloomington *Drosophila* Stock Center: UAS*-tara-*RNAi (JF01421, # 31634), UAS*-dicer2* (*dcr2,* #24650), *tara*^1^ (#6403), UAS*-vri* (#78077), *elav-Gal4* (#458)*, nos-Cas9* (#54591), *tim-Gal4* (#7126), *Tim*-UAS-Gal4 (TUG; #80941), *Pdf-Gal4* (#80939), and the background control line *iso31* (*w*^1118^, #3605). The UAS*-tara-*RNAi (6889R-1, #NM_080255.2) line was obtained from the National Institute of Genetics, Japan. UAS-*Pdp1* and *tara*^s132^ were previously described (Zheng et al., 2009; Afonso et al., 2015b). UAS*-dcr2* was combined with two RNAi lines (JF01421 and 6889R-1) to enhance the efficiency of *tara* knockdown. TARA and its mammalian homologs have four conserved domains (Calgaro et al., 2002): CycA binding homology (CAB), SERTA, PHD-Bromo binding (PBB), and C-terminal (CT) . UAS-*tara* (containing the full-length *tara-B* coding sequence) and *tara* domain deletion (ΔCAB, ΔSERTA, ΔPBB, and ΔCT) lines were generated in this study. All fly lines were outcrossed with *iso31* for at least five generations, except for the UAS*-tara*-RNAi and *nos-Cas9* lines. At the final outcrossing step, paired control and mutant lines were derived from the same heterozygous mutant parents.

### Circadian assays

To conduct the circadian assays, 3 to 5-day-old flies entrained to an LD cycle at 25°C for at least 3 days were placed individually in small glass tubes containing 5% sucrose and 2% agar. The *Drosophila* Activity Monitoring System (Trikinetics) was used to record circadian locomotor activity for ∼6.5 days in constant darkness (DD) and ∼2.5 days in LD. Data for 6 days in DD and 2 days in LD were analyzed for free-running rhythms and anticipatory behavior, respectively. We employed the Faas software (M. Boudinot and F. Rouyer, Institute of Neurosciences Paris-Saclay, France) to calculate the circadian period length and rhythm strength of each fly assayed in DD. The software analyzes activity data collected in 30-min bins using χ^2^ periodogram analysis to determine the period length and rhythm strength. Rhythm strength is calculated as the difference between the peak χ^2^ value and the expected χ^2^ value at chance (p = 0.05). Based on the rhythm strength value, each fly was classified as arrhythmic (<25), weakly rhythmic (25-50), or rhythmic (>50). All flies were included in the statistical analysis of rhythm strength, while the analysis of period length excluded arrhythmic flies. Including rhythm strengths of arrhythmic flies provides a single measure of rhythmicity for a group of flies instead of two measures (% rhythmic flies and the average rhythm strength of rhythmic flies), simplifying statistical comparisons. The anticipation index, defined as the ratio between the total activity counts during a 3-h period immediately before a light-dark transition and a 6-h period before the transition, was calculated using Excel. Activity counts averaged over 2 days were used for anticipation index calculations.

### Immunohistochemistry

Following dissection, adult brains were fixed in 4% paraformaldehyde for 30 min at room temperature. The samples were blocked in 1% bovine serum albumin (BSA) for PER and PDF antibody staining. The primary and secondary antibodies were diluted in 0.1% BSA. The primary antibodies utilized were anti-PER (rabbit 13.1 from the Sehgal lab, University of Pennsylvania) at a dilution of 1:8000 and mouse anti-PDF (mouse C7s from the Developmental Studies Hybridoma Bank) at a dilution of 1:2000. The secondary antibodies, Alexa Fluor 647 goat anti-rabbit and Alexa Fluor 488 goat anti-mouse (Invitrogen) were used at a dilution of 1:1000. Both primary and secondary antibodies were incubated overnight at 4°C, and images were captured using a Leica SP8 confocal microscope.

### Plasmid constructs for cell-based assays

*Actin-Clk, Actin-per, Actin*-*Gal4, per-enhanced luciferase* (*per-luc*) (Darlington et al., 1998), and pUAST-V5 vector were obtained from the Sehgal lab (University of Pennsylvania). *Clk-luc* (∼3 kb *Clk* promoter region fused to a *luciferase* cDNA, UAS*-vri, and* CMV*-vri::VP16* (Cyran et al., 2003) were obtained from the Blau lab (New York University). UAS*-Pdp1* was previously described (Zheng et al., 2009). CMV-FLAG::*hTRIP-Br1* and CMV-FLAG::*hTRIP-Br2* were a gift from Dr. Jit Kong Cheong (National University of Singapore). *Actin-Renilla luciferease* (*rluc*) and CMV*-rluc* (Promega) were used to control for transfection efficiency.

A full-length, untagged UAS-*tara* construct was generated by PCR-amplifying the entire *tara*-B coding region from *tara* cDNA and inserting it into the pUAST vector (Brand and Perrimon, 1993). UAS-HA*::tara* constructs were generated by inserting the *tara-B* coding region, amplified from the untagged UAS-*tara* construct, into the pTHW Gateway vector (*Drosophila* Genomics Resource Center) containing the UAST promoter and N-terminal 3xHA tag. The TARA-B residues removed in the deletion constructs are as follows: ΔCAB:1-26, ΔSERTA: 432-479, ΔPBB: 722-739, and ΔCT: 893-912. pUAST-attB-V5::*tara* was generated by inserting the *tara-B* coding region into the pUAST-attB-V5 vector, which is derived from the pUAST-attB vector (*Drosophila* Genomics Resource Center) for N-terminal V5 tagging. CMV-*tara* was generated by digesting UAS-*tara* with BglII and XbaI and inserting the *tara-B* coding sequence into the BamHI and XbaI sites in the pCDNA3 vector (Invitrogen). UAS-*Pdp1*::V5 was generated by digesting UAS-*Pdp1* (Zheng et al., 2009) with NotI and XhoI, and inserting the *Pdp1* coding sequence into the corresponding sites in the pUAST-V5 vector, which is derived from pUAST for C-terminal V5 tagging. UAS-*vri*::MYC was generated by inserting the *vri* coding sequence, amplified from a UAS-*vri* construct (Cyran et al., 2003), into the pTWM Gateway vector (*Drosophila* Genomics Resource Center). The constructs were validated by sequencing (GENEWIZ).

### Luciferase-based transcription assay

*Drosophila* S2 cells were cultured at 25°C in Schneider’s Medium (Gibco), while HEK293 cells were cultured at 37°C in DMEM Medium (Gibco). Both media were supplemented with 10% heat-inactivated fetal bovine serum and 1% penicillin/streptomycin. Approximately 24 h prior to transient transfection, 48-well plates were seeded with ∼1.5 x 10^5^ *Drosophila* S2 or HEK293 cells and maintained at their respective temperatures (25°C or 37°C). Transfections were performed with Effectene Reagent (Qiagen) per the manufacturer’s protocol. Transcriptional activity was determined using the Dual Luciferase Assay Kit (Promega). Approximately 48 h after transfection, cells were lysed in 1X Passive Lysis Buffer (PLB), and luciferase activity was measured using the Luciferase Assay Reagent in Cytation 5 Luminometer (Biotek) following the manufacturer’s instructions.

For the PDP1- and VRI-dependent transcriptional assay, the following constructs were transfected: *Clk-luc* (10 ng), *Actin-rluc* (10 ng), *Actin-Gal4* (70 ng), UAS*-Pdp1*::V5 (10 ng), UAS*-vri*::MYC (35 ng), and UAS*-*HA*::tara* (1 ng, 5 ng, or 15 ng). An empty pUAST::V5 vector was included to ensure an equal amount of total DNA (150 ng) in all conditions. For the CLK-dependent transcriptional assays, the following constructs were used: *per*-*luc* (20 ng), *Actin-rluc* (5 ng), *Actin-Gal4* (20 ng), *Actin-Clk* (1 ng), UAS-HA::*tara* (10 ng or 30 ng). Empty *pActin* and pUAST::V5 vectors were used to control for the total DNA amounts (150 ng). For the PER-dependent transcriptional assay, the following constructs were transfected: *per*-*luc* (20 ng), *Actin-rluc* (15 ng), *Actin-Gal4* (20 ng), *Actin-Clk* (1 ng), *Actin-per* (2 ng, 5 ng, or 15 ng), and UAS-HA*::tara* (30 ng). Empty *pActin* and pUAST::V5 vectors were used to control for the total DNA amounts (150 ng). For the VRI::VP16-dependent transcriptional assay, the following constructs were transfected: *Clk-luc* (10 ng), CMV*-rluc* (10 ng), *Actin-Gal4* (70 ng), CMV*-vri::VP16* (5 ng), CMV*-tara* (10ng or 30ng), CMV-HA::ΔSERTA (30 ng), CMV-FLAG*::hTRIP-Br1* (30 ng), CMV-FLAG*::hTRIP-Br2* (30 ng), and the pCDNA3 vector to control for the total DNA amount (125 ng).

### Western analysis and co-immunoprecipitation (co-IP)

To examine the expression levels of transfected proteins, *Drosophila* S2 cell lysates were collected approximately 48 h after transfection. Cells were lysed in 1X PLB to release cytoplasmic proteins, while the insoluble fractions were treated with a sample buffer containing sodium dodecyl sulfate (SDS) to release nuclear proteins. For the co-IP assay, ∼3.2x10^6^ S2 cells were seeded in each well of a 6-well plate and transfected ∼24 h after seeding. Around 48 h after transfection, cells were harvested and lysed in 1X RIPA buffer supplemented with Protease Inhibitor Cocktail (Sigma). The cell lysates were pre-cleaned with Dynabeads Protein G (Invitrogen) to avoid non-specific binding. The pre-cleaned lysates were incubated with anti-V5-conjugated magnetic beads (Medical & Biological Laboratories) overnight at 4°C. To elute the protein complexes bound to the beads, 2X SDS with 5% β-mercaptoethanol was used. Western blot analysis was performed as described (Wu et al., 2010). Antibodies against TARA (TJR51, 1:1000; (Afonso et al., 2015b), HA (Covance, 1:1000), PDP1 (1:1000; Cyran et al., 2003), β-ACTIN (Abcam, 1:8000), and Histone H3 (Abcam, 1:3000) were used.

### Generation of tara mutants with domain deletions and a UAS-tara transgenic line

To identify an optimal protospacer adjacent motif (PAM) sequence within each domain, we utilized the flyCRISPR target finder (https://flycrispr.org/target-finder). The following PAM (shown in bold) and their corresponding flanking sequences were identified, with the numbers indicating the nucleotide positions in the *tara-b* transcript: CAB domain: 552-574 – **CCA** AGC GGA AGC ATG AGC TGA CC; SERTA domain: 1781-1802 – G CCA ACA GGA GTT GCT ACA **AGG**; PBB domain: 2672-2680 – TA CTA GCG ATT CGG GTT ATG **CGG**; CT domain: 3186-3208 – **CCC** GCC TGC ACG ACA ACG AAC TG. To introduce the domain-specific protospacer sequences into the pCFD4 vector (Addgene), we employed the ligation-independent Gibson Assembly method (NEBuilder, New England BioLabs) according to the manufacturer’s protocol. Transgenic flies carrying domain-specific gRNAs were generated using the standard embryo injection technique (Rainbow Transgenics). Domain-specific deletions were generated by crossing fly lines carrying the domain specific gRNAs with a *nos*-*Cas9* fly line. Mutant flies were identified by sequencing the genomic DNA (GENEWIZ). Additionally, a transgenic fly line carrying the untagged UAS-*tara* construct was generated using the standard embryo injection technique (Rainbow Transgenics). We confirmed that the UAS-*tara* transgene can increase TARA expression when driven by a salivary gland driver by immunostaining with an antibody against TARA (TJR51; Afonso et al., 2015b). We used salivary glands because TARA overexpression using *elav*-Gal4 led to lethality, and the TARA antibody could not detect TARA expression in s-LN_v_s when overexpressed using *Pdf*-Gal4.

### RNA extraction and quantitative real-time PCR (qPCR)

Total RNA was extracted from 30 male fly heads using 1 mL of TRIzol Reagent (Ambion). We employed 100 µL of 1-Bromo-3-Chloropropane (Molecular Research Center, Inc.) to separate the phases, and RNA precipitation was achieved by adding 500 µL of 2-propanol. After treatment with 2 µL of DNaseI, cDNA was amplified from 1 µg of RNA using a high-capacity cDNA Reverse Transcription Kit (Applied Biosystems). For qPCR, we used SYBR Green PCR Master Mix (Applied Biosystems) and the 7500 Real-Time PCR System (Applied Biosystems). 3 technical replicates of 6 independent biological samples were assayed for each genotype. The following primers were used for *Pdf*: 5’-CCT TGT GCT TCT GGC CAT TT-3’ and 5’-GTC GAG GAG ATC CCG ATT GT-3’. *actin* mRNA was used for normalization.

### Experimental design and statistical analyses

Throughout the study, we employed a between-subject design. Immunohistochemistry and behavioral experiments were repeated at least twice using flies from independent crosses, while cell-based experiments were conducted at least three times. Statistical analyses were performed using GraphPad Prism 9. The results of statistical tests are presented in Table 1. All datasets were assessed for normality using Kolmogorov Smirnov tests, and we employed non-parametric tests when the normality tests failed. When performing parametric tests, we used tests that do not assume homogeneity of the variances. Comparisons between two groups were performed using non-parametric Mann-Whitney or parametric Welch’s t-tests. Comparisons among multiple groups were performed using non-parametric Kruskal-Wallis or parametric one-way Brown-Forsythe and Welch ANOVA tests. For experiments involving two factors, two-way ANOVA tests were employed. Post hoc comparisons employed Dunn’s, Tukey’s, or Sidak’s multiple comparisons tests, as indicated in figure legends and Table 1.

**Table 1:**
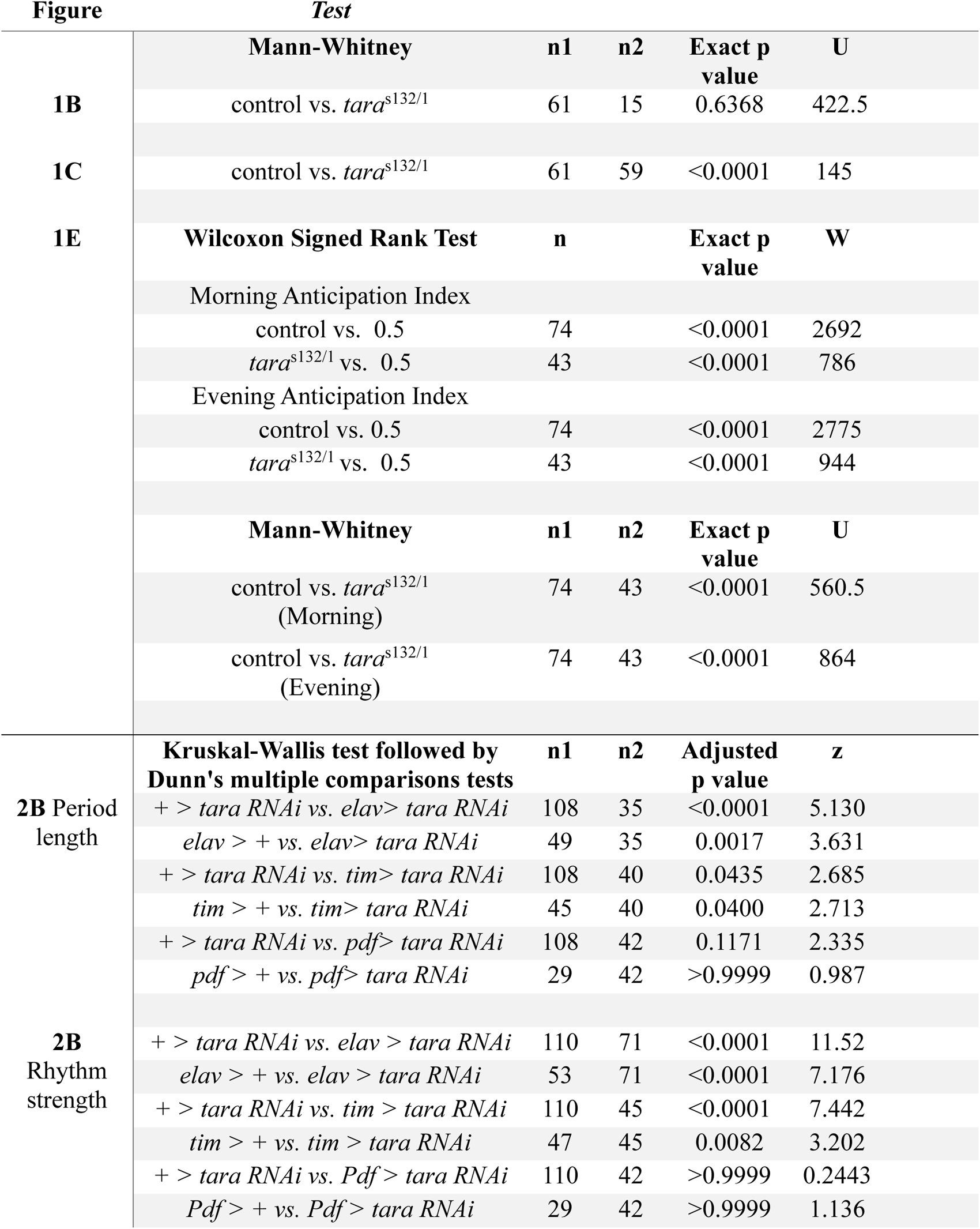
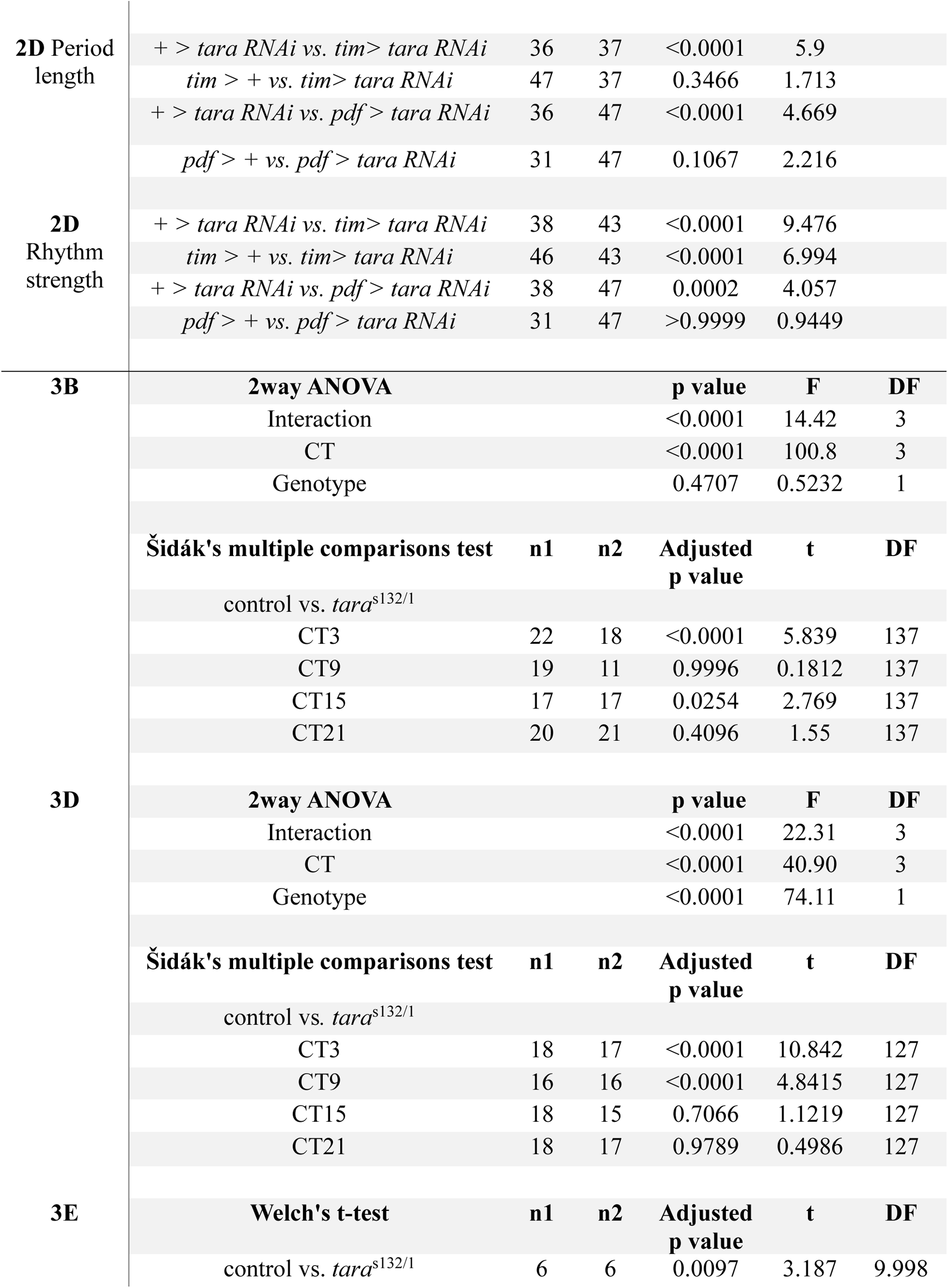
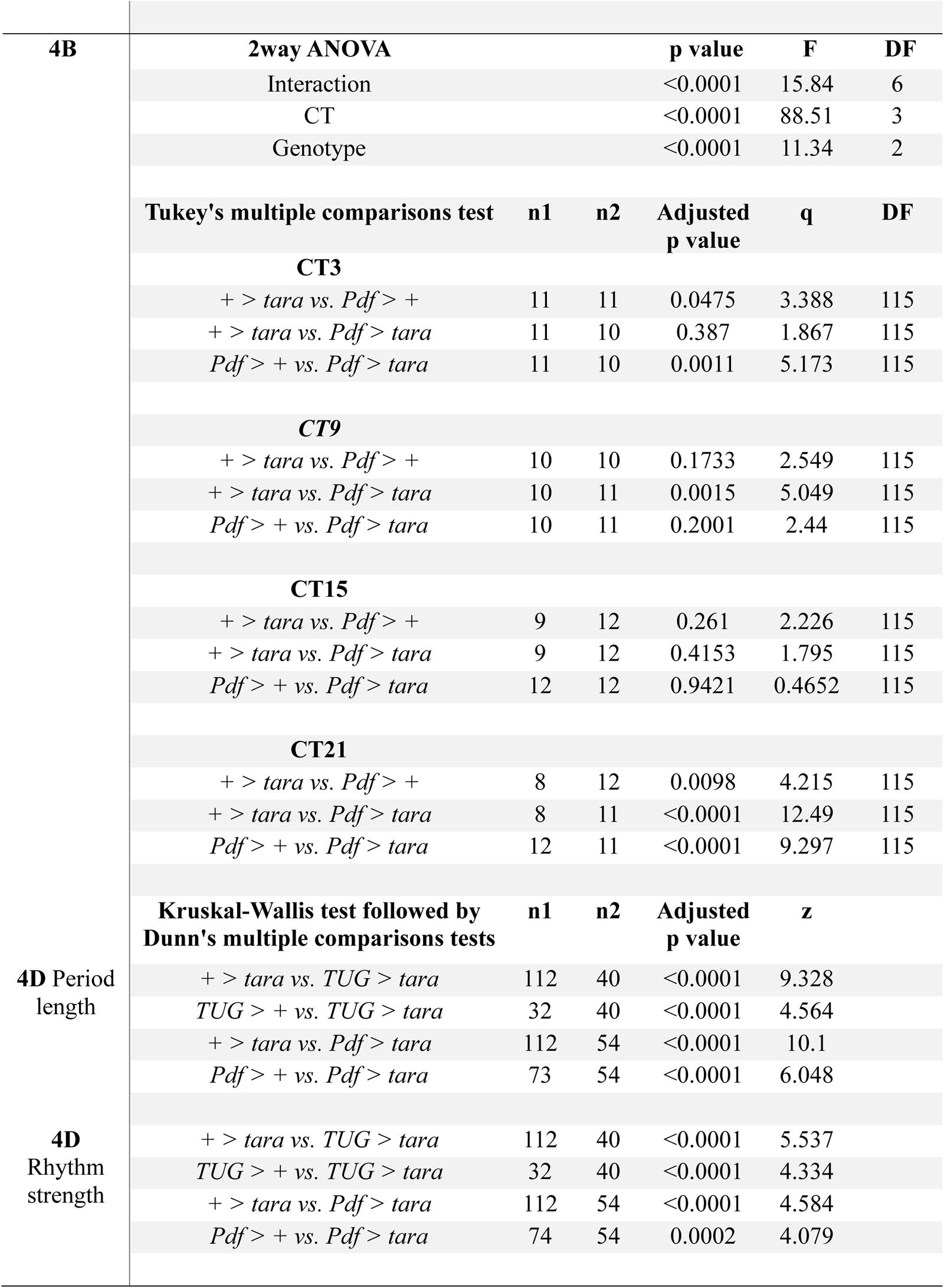
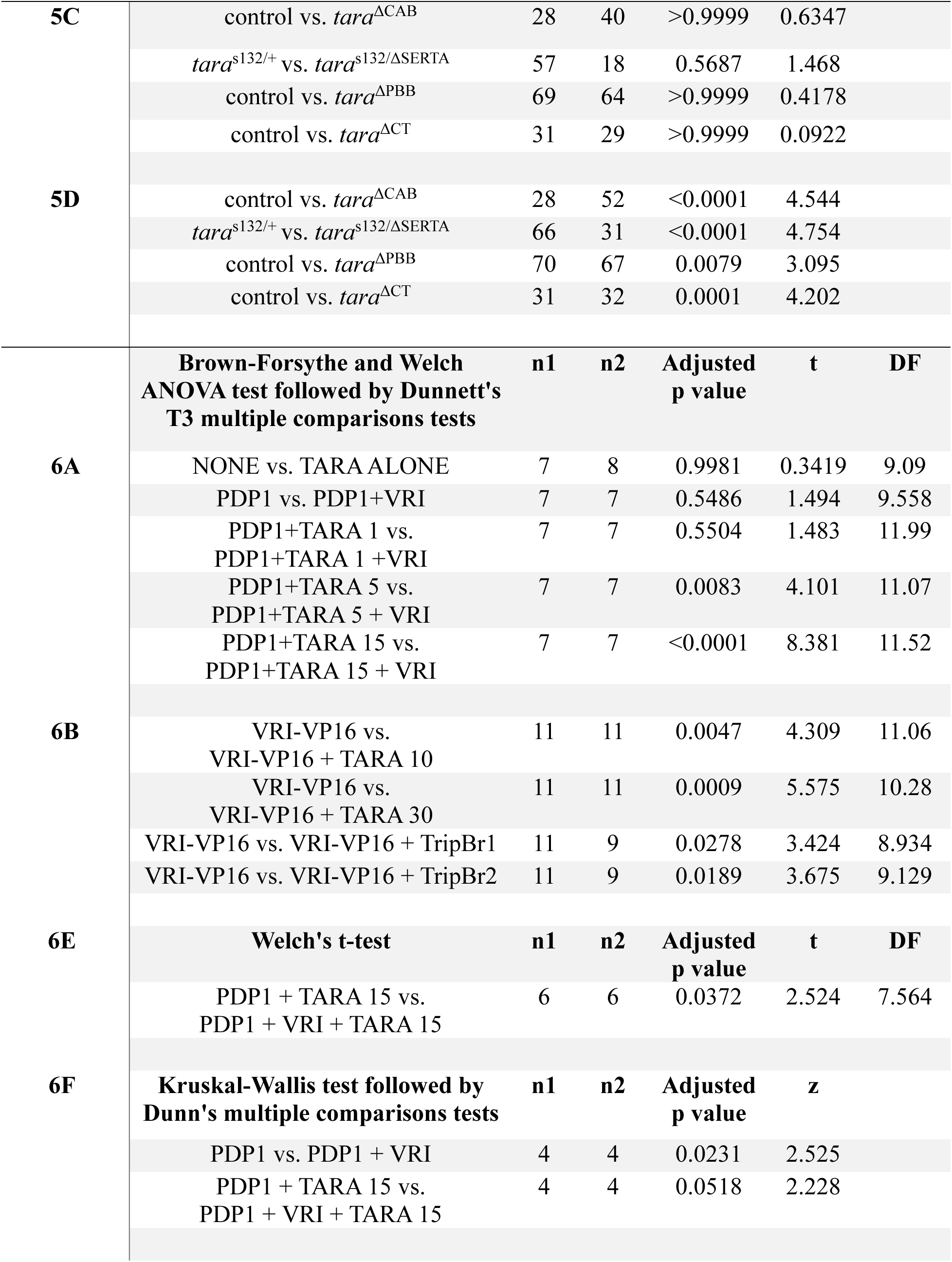
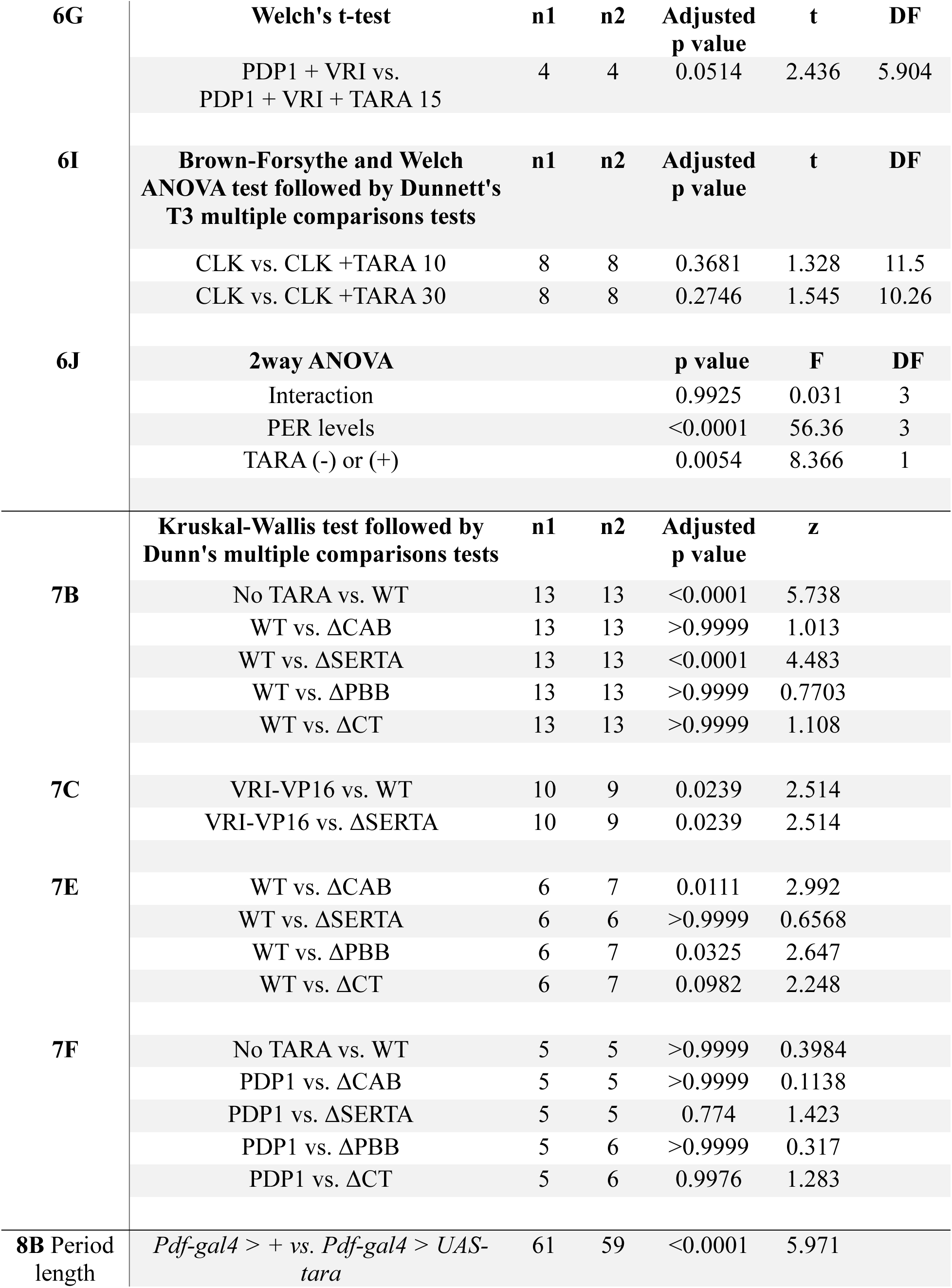
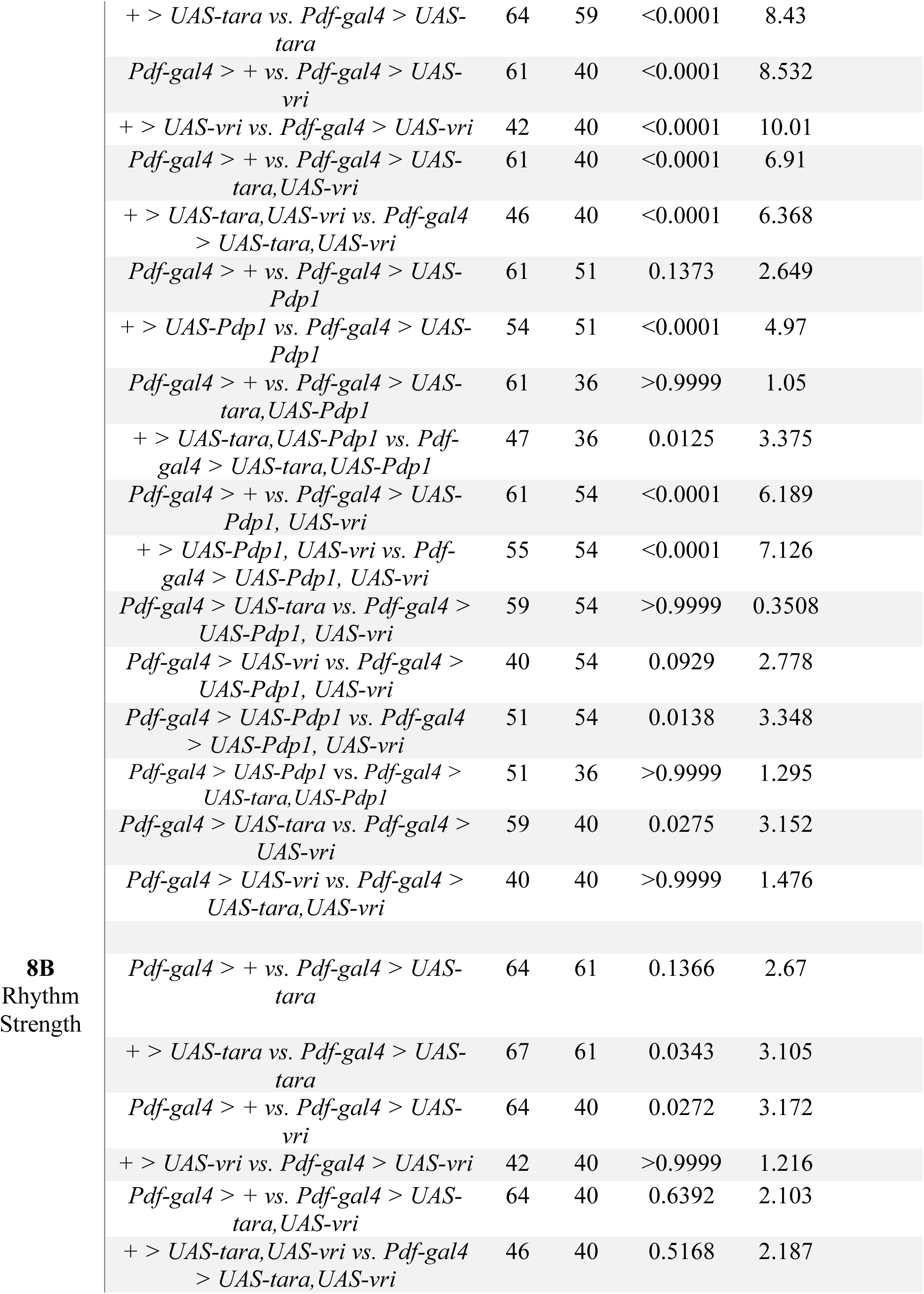
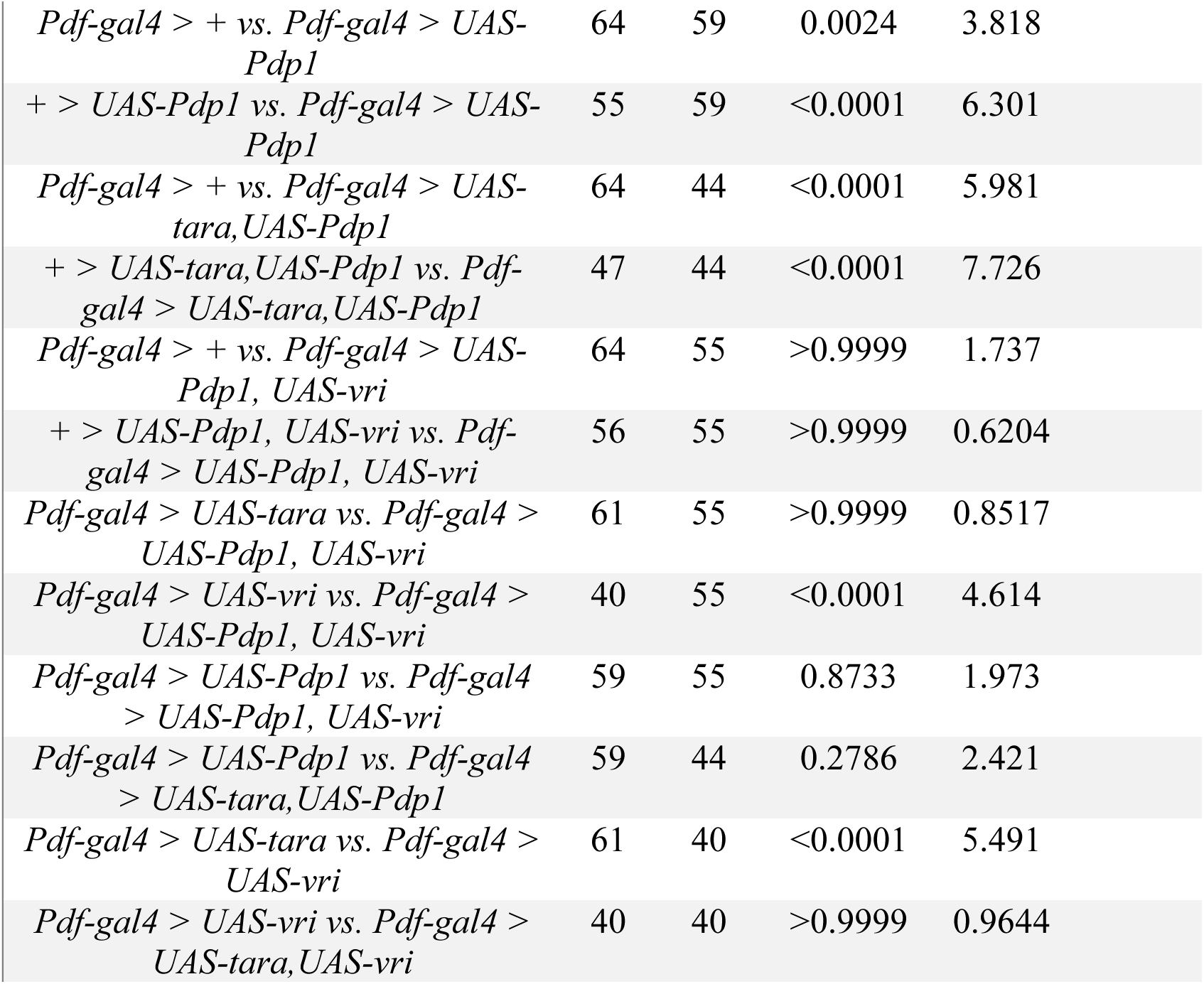
Statistical Analysis grouped by figure number, figure panel, and statistical tests.

## Results

### TARA functions in clock neurons to promote rhythmic behavior

We previously showed that *tara* mutants display reduced sleep in LD, DD, and constant light (LL) conditions, suggesting that the short-sleep phenotype is independent of the circadian clock and light conditions (Afonso et al., 2015b). Additionally, the study found that *tara* mutants exhibit reduced rhythmicity. We confirmed markedly reduced rhythmicity in *tara*^s132/1^ mutants in DD (Tables 1-2; Fig. 1A-C). Activity patterns in LD conditions revealed that *tara* mutants exhibited elevated activity levels compared to controls, especially at night (Fig. 1D). To examine rhythmic behavior in LD, we analyzed morning and evening anticipation, i.e., the increase in locomotor activity before lights on and off, respectively. We found that *tara* mutants exhibited significant morning and evening anticipation (Fig. 1D,E). However, *tara* mutants had lower anticipation indexes than control flies, which may result from their higher overall activity levels. These findings demonstrate that reduced *tara* leads to decreased rhythmicity in locomotor behavior in LD as well as DD.

**Figure 1.**
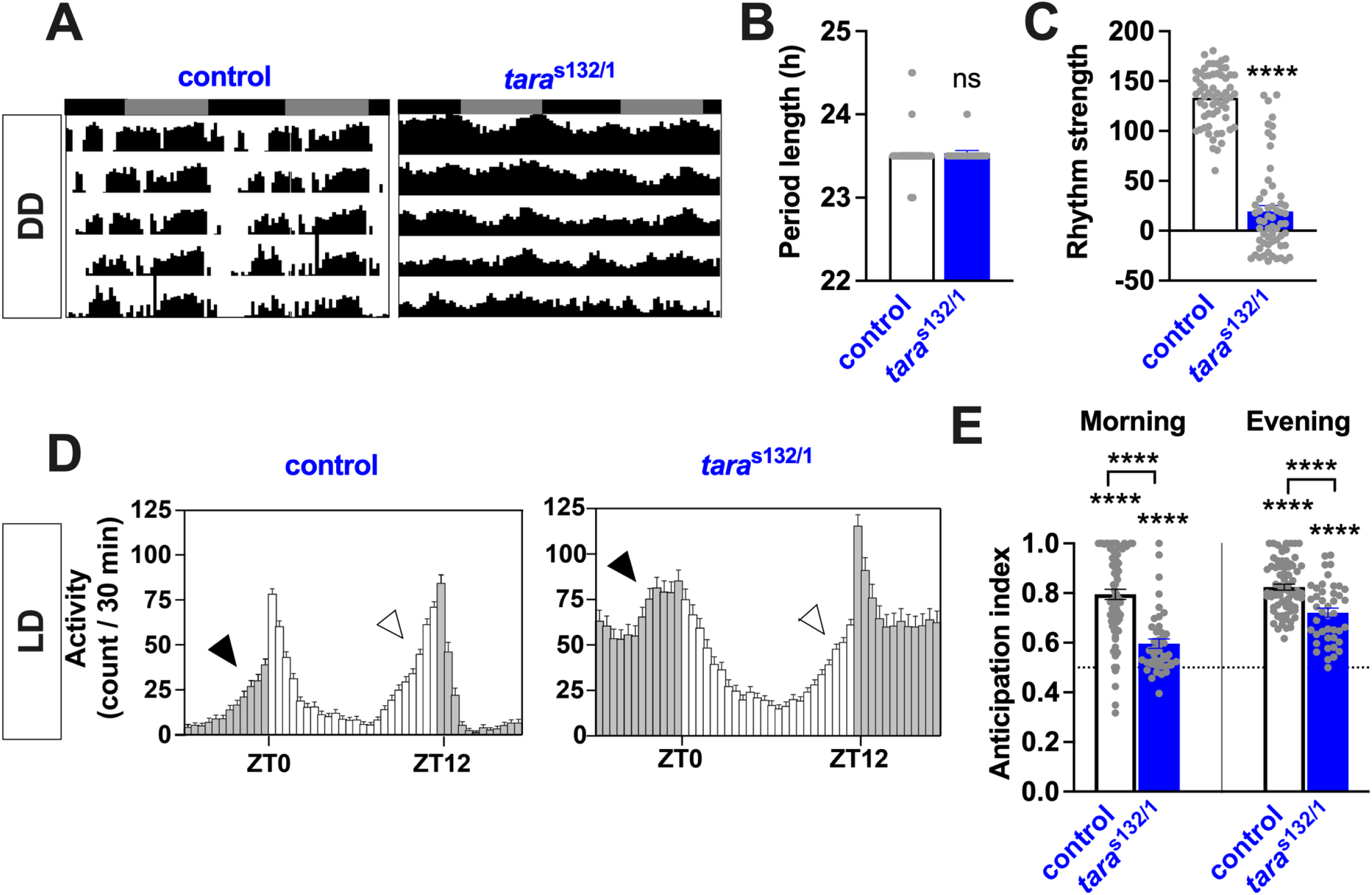
*tara* mutants have impaired locomotor rhythms and reduced morning and evening anticipatory behavior. **(A)** Representative circadian actograms of control and *tara*^s132/1^ male flies. Mutants were outcrossed to *w*^-^ (iso31) background 5 times, and paired control and mutant lines were derived from the same heterozygous mutant parents. Control flies for *tara*^s132/1^ transheterozygous mutants were generate by crossing the control lines for *tara*^s132^ and *tara*^1^. Flies were entrained at least 3 days in 12 h:12 h LD before being monitored in DD. In this figure and subsequent figures, actograms present 6 days of activity in DD, and the gray and black bars above the actogram indicate subjective day and night, respectively. **(B-C)** Period length (B) and rhythm strength (C) of indicated genotypes. Mean ± SEM is shown. N = 15-61 for period length; n = 59-61 for rhythm strength. **(D)** Activity profiles of control and *tara*^s132/1^ males under 12 h:12 h LD conditions. Activity counts averaged over 2 days are presented. Filled and open arrowheads point to morning and evening anticipation, respectively. **(E)** Morning and evening anticipation index for control and *tara*^s132/1^ male flies. The anticipation index is defined as the ratio of the activity counts within 3 h immediately before a light-dark transition and those over 6 h before the transition. Activity counts averaged over 2 days were used for anticipation index calculations. N = 43-74. Bars represent mean anticipation index ± SEM. Statistics above the individual bars indicate whether the anticipation index of each genotype is significantly different from 0.5 (dotted line), which represents no anticipation, while those above the brackets spanning pairs of bars indicate whether the two genotypes are different from each other. ****p < 0.0001, ns: not significant; Mann-Whitney test (B,C) and Welch’s t-test (E).

**Table 2.** Circadian locomotor rhythm phenotypes in DD.

| Genotype | N | % R | % WR | % AR | tau (h) ± SEM | Power ± SEM |
| --- | --- | --- | --- | --- | --- | --- |
| ***tara* mutant** |  |  |  |  |  |  |
| control | 61 | 100 | 0 | 0 | 23.45 ± 0.02 | 133.30 ± 3.53 |
| *tara*<sup>s132/1</sup> | 59 | 25.4 | 13.6 | 61 | 23.50 ± 0.00 | 19.42 ± 5.93 |
| **Knockdown** |  |  |  |  |  |  |
| <u>Females</u> |  |  |  |  |  |  |
| + > *tara* RNAi | 110 | 98.1 | 0.9 | 0.9 | 24.58 ± 0.04 | 115.8 ± 2.38 |
| *elav* > + | 53 | 88.7 | 3.8 | 7.5 | 24.47 ± 0.08 | 98.3 ± 5.14 |
| *elav* > *tara* RNAi | 71 | 26.8 | 21.1 | 52.1 | 24.03 ± 0.09 | 30.6 ± 4.86 |
| *tim* > + | 47 | 83.1 | 14.8 | 2.1 | 24.64 ± 0.05 | 91.7 ± 5.05 |
| *tim* > *tara* RNAi | 45 | 60 | 28.9 | 11.1 | 24.36 ± 0.07 | 62.8 ± 4.90 |
| *Pdf* > + | 29 | 100 | 0 | 0 | 24.86 ± 0.05 | 106.0 ± 4.57 |
| *Pdf* > *tara* RNAi | 42 | 100 | 0 | 0 | 24.79 ± 0.06 | 114.6 ± 3.32 |
| <u>Males</u> |  |  |  |  |  |  |
| + > *tara* RNAi | 38 | 100 | 0 | 0 | 23.66 ± 0.05 | 153.6 ± 3.40 |
| *tim* > + | 47 | 100 | 0 | 0 | 24.39 ± 0.05 | 135.6 ± 3.23 |
| *tim* > *tara* RNAi | 43 | 72 | 14 | 14 | 24.26 ± 0.08 | 69.38 ± 5.90 |
| *Pdf* > + | 31 | 100 | 0 | 0 | 24.31 ± 0.06 | 137.6 ± 2.98 |
| *Pdf* > *tara* RNAi | 47 | 100 | 0 | 0 | 24.09 ± 0.05 | 131.6 ± 3.04 |
| **Overexpression** |  |  |  |  |  |  |
| <u>Males</u> |  |  |  |  |  |  |
| + > *tara* | 112 | 100 | 0 | 0 | 23.75 ± 0.04 | 133.4 ± 2.85 |
| *TUG* > + | 32 | 100 | 0 | 0 | 24.13 ± 0.07 | 134.6 ± 3.08 |
| *TUG* > *tara* | 40 | 100 | 0 | 0 | 24.81 ± 0.09 | 105.6 ± 4.37 |
| *Pdf* > + | 74 | 98.6 | 0 | 1.4 | 24.10 ± 0.05 | 132.6 ± 3.03 |
| *Pdf* > *tara* | 54 | 100 | 0 | 0 | 24.73 ± 0.04 | 116.3 ± 2.44 |

N: number of flies; R: rhythmic; WR: weakly rhythmic; AR: arrhythmic.
tau: free-running period; Power: a measure of rhythm strength

We confirmed that the reduced rhythmicity maps to the *tara* locus using RNAi-mediated *tara* knockdown in all neurons using *elav*-*Gal4* (Table 2; Fig. 2 A,B). The *Drosophila* brain contains ∼150 circadian neurons in several clusters. A cluster of clock neurons expressing the Pigment Dispensing Factor (PDF) neuropeptide, called small ventral lateral neurons (s-LN_v_s), is essential for free-running rhythms in DD (Renn et al., 1999). To examine where TARA functions to promote rhythmic behavior, we employed *tim*-*Gal4*, which targets all clock neurons, and *Pdf-Gal4*, which targets the PDF-expressing LN_v_s. Male flies are typically used for assaying locomotor rhythms because they display more robust and consistent rhythms than females. However, because pan-neuronal *tara* knockdown led to lethality in males, female flies were used in the pan-neuronal knockdown experiment. Both males and females were assayed in experiments using *tim*-*Gal4* and *Pdf*-*Gal4*.

**Figure 2.**
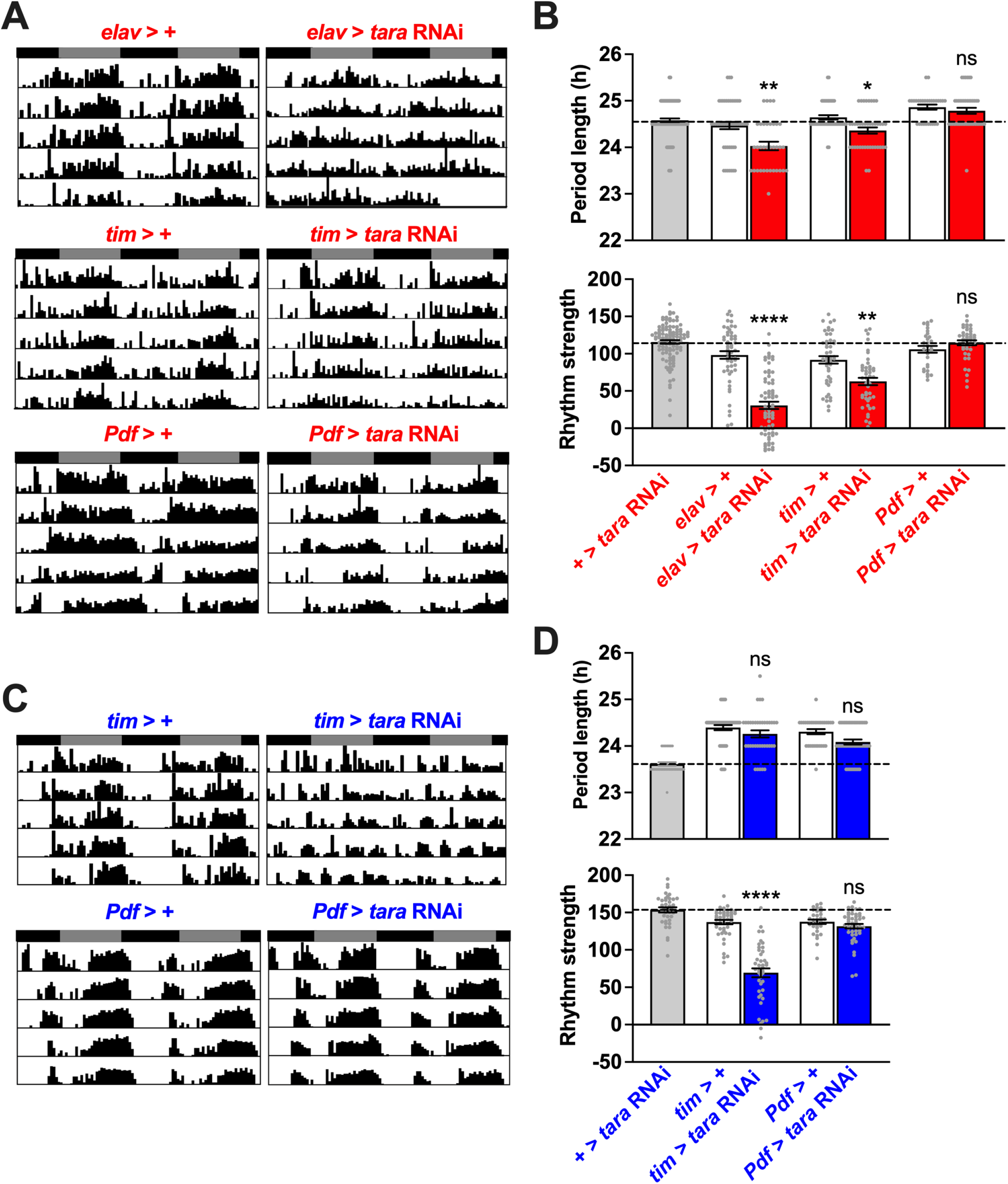
Pan-neuronal and clock-neuron-specific knockdown of *tara* degrades rhythm strength and can shorten the circadian period. **(A)** Representative circadian actograms of females of the following genotypes in DD: pan-neuronal knockdown (*elav* > *tara* RNAi), clock-neuron-specific knockdown (*tim* > *tara* RNAi), LN_v_-specific knockdown (*Pdf* > *tara* RNAi), and their respective *Gal4* driver controls (*elav* > +, *tim* > +, and *Pdf* > +). **(B)** Period length (top) and rhythm strength (bottom) of females of indicated genotypes. N = 29-108 for period length; n = 29-110 for rhythm strength. Red labels indicate female flies, while blue labels indicate male flies. The horizontal dotted line represents the mean period length (top) and rhythm strength (bottom) of the common control (+ > *tara* RNAi). **(C)** Representative circadian actograms of males of indicated genotypes in DD. **(D)** Period length (top) and rhythm strength (bottom) of males of indicated genotypes. N = 31-47 for period length and rhythm strength. Mean ± SEM is shown. ****p < 0.0001, ns: not significant; Kruskal-Wallis test followed by Dunn’s multiple comparisons test relative to both parental controls (B and D).

Pan-neuronal *tara* knockdown using *elav-Gal4* and knockdown in all clock neurons using *tim-Gal4* resulted in a small but significant reduction in period length in females (Table 2, Fig. 2). In contrast, *tara* knockdown using *Pdf*-Gal4 in females and males or *tim*-Gal4 in males did not result in shortened periods. In the former cases, the Gal4 and UAS controls had similar period lengths, while in the latter cases, the Gal4 control flies showed substantially longer periods than the UAS control flies. The longer periods of the Gal4 control flies may have made it difficult to detect an underlying period-shortening effect of *tara* knockdown in PDF+ neurons. In both males and females, knockdown of *tara* in all clock neurons using *tim*-*Gal4* resulted in a significant reduction of rhythm strength (Table 2, Fig. 2). In contrast, restricted *tara* knockdown in PDF-expressing LN_v_s did not significantly affect rhythmicity. This finding suggests that TARA functions in multiple clock neuron clusters to regulate rhythm strength. Comparing female data with *elav*-*Gal4* vs. *tim*-*Gal4*, we observed that pan-neuronal knockdown of *tara* had a more substantial effect on rhythm strength than knockdown in clock neurons (Table 2, Fig 2A,B), suggesting TARA also functions in non-clock neurons to promote rhythmic locomotor activity. Our results suggest that TARA promotes rhythmic locomotion in clock and non-clock neurons and may modulate the pace of the clock.

### TARA modulates the speed of the circadian clock

We previously showed that TARA is expressed in s-LN_v_s (Afonso et al., 2015a). To investigate whether TARA influences molecular clock oscillation, we examined PER protein expression in s-LN_v_s on Day 3 in DD. Although ∼75-80% of *tara*^s132/1^ mutants are arrhythmic or weakly rhythmic (Afonso et al., 2015b; Table 2), they displayed robust daily cycling of PER levels in DD (Fig. 3A,B), suggesting that TARA regulates locomotor rhythms downstream of the s-LNv molecular clock. TARA may regulate the output of the s-LN_v_s or other clock neurons. Compared to control brains, *tara* mutant brains exhibited a significantly lower PER signal at CT 3 and a substantially higher PER signal at CT 15 (Fig. 3A,B), a pattern consistent with a faster molecular clock. Because only 4 timepoints over a single cycle were examined, these findings are insufficient to show that the molecular clock is running faster in *tara* mutants. However, considering the significantly shorter periods in female flies in which *tara* was knocked down pan-neuronally or in all clock neurons, TARA likely plays a role in modulating the speed of the clock. *tara*^s132/1^ mutants, which have the most severe reduction in *tara* levels among viable *tara* mutants, did not exhibit shorter periods in locomotor behavior (Table 2, Fig. 1A-C; Afonso et al., 2015b). This may be because the more strongly affected mutants were arrhythmic, leaving only the less severely affected mutants for period length measurements.

**Figure 3.**
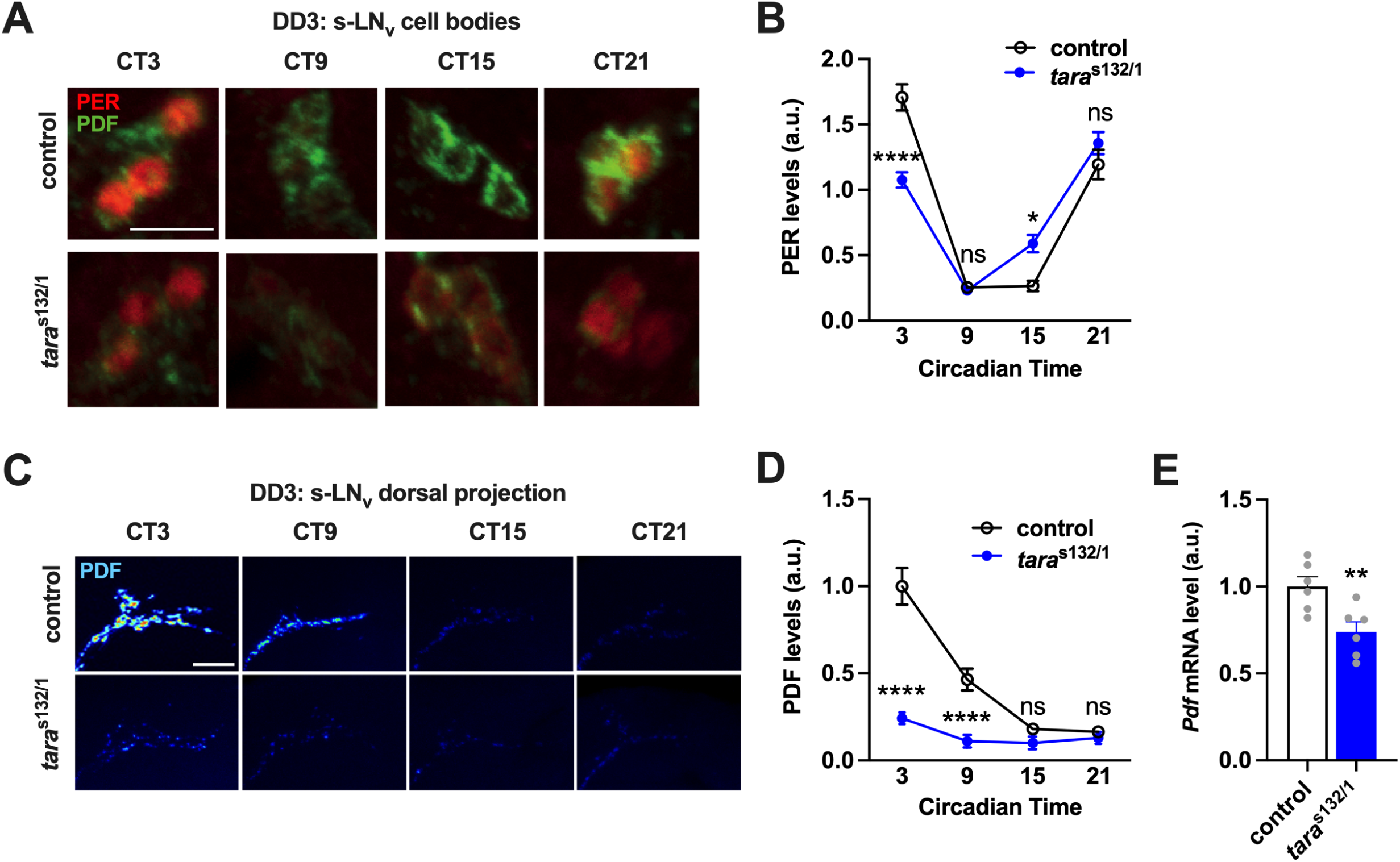
PER cycling and PDF accumulation are altered in the s-LN_v_ of *tara* mutants. (A) **(A)** Representative expression patterns of PER and PDF in the s-LN_v_s of control and *tara*^s132/1^ males at the indicated circadian times on the 3^rd^ day in DD. The scale bar is 10 μm. **(B)** Quantification of PER levels in s-LN_v_s for control and *tara*^s132/1^ males (n = 11-19). **(C)** Representative PDF expression patterns in the dorsal projections of s-LN_v_s at the indicated circadian times on the 3^rd^ day in DD in control and *tara*^s132/1^ males. Signal intensity is indicated using the “royal” lookup table, where black represents the minimum intensity, and white represents the maximum intensity. The scale bar is 20 μm. **(D)** Quantification of PDF levels in the dorsal projection of s-LN_v_s for control and *tara*^s132/1^ males (n = 15-18). **(E)** Quantification of *Pdf* mRNA levels from whole head extracts for control and *tara*^s132/1^ males. Six independent biological samples collected at CT15 on DD3, each from 30 fly heads, were assayed. Mean ± SEM is shown. ****p < 0.0001, ns: not significant; two-way ANOVA followed by Sidak’s multiple comparisons tests (B and D); Welch’s t-test (E).

In addition to the PER cycling changes, we observed that PDF levels at the dorsal projections of the s-LN_v_s were considerably lower in *tara* mutants than in control flies on Day 3 in DD (Fig. 3C,D). We also found a significant reduction in the total levels of *Pdf* mRNA in *tara* mutants compared to the controls (Fig. 3E). However, the reduction in PDF protein (∼75%) was more pronounced than that of the *Pdf* transcripts (∼25%), suggesting that TARA impacts PDF levels at both the transcriptional and post-transcriptional levels. Lower PDF protein levels in *tara* mutants may be due to reduced synthsis, increased degradation, or increased release. *Pdf* null mutants display decreased rhythmicity and shorter periods (Renn et al., 1999). Reduced PDF levels in *tara* mutants may contribute to their behavioral and molecular phenotypes, even though it is unlikely to be their sole reason.

To further examine the effect of TARA levels on the speed of the circadian clock, we examined whether flies overexpressing TARA exhibit the opposite pattern of PER cycling as *tara* mutants. Indeed, TARA overexpression in PDF-expressing (PDF+) neurons led to a significantly lower PER signal at CT 21 on Day 3 in DD, consistent with a slower molecular clock (Fig. 4A,B). The small reduction in overall PER levels in *tara* overexpressing flies relative to controls could be because the timepoints we sampled did not include the peak PER expression in *tara* overexpression flies. Nevertheless, flies overexpressing TARA in all clock neurons or PDF+ neurons exhibited a significantly longer circadian period than control flies (Table 2, Fig. 4 C,D), supporting TARA’s role in regulating the clock speed. Collectively, our data suggest that *tara* plays a dual role in setting the pace of the molecular clock and regulating locomotor rhythmicity downstream of the s-LNv molecular clock.

**Figure 4.**
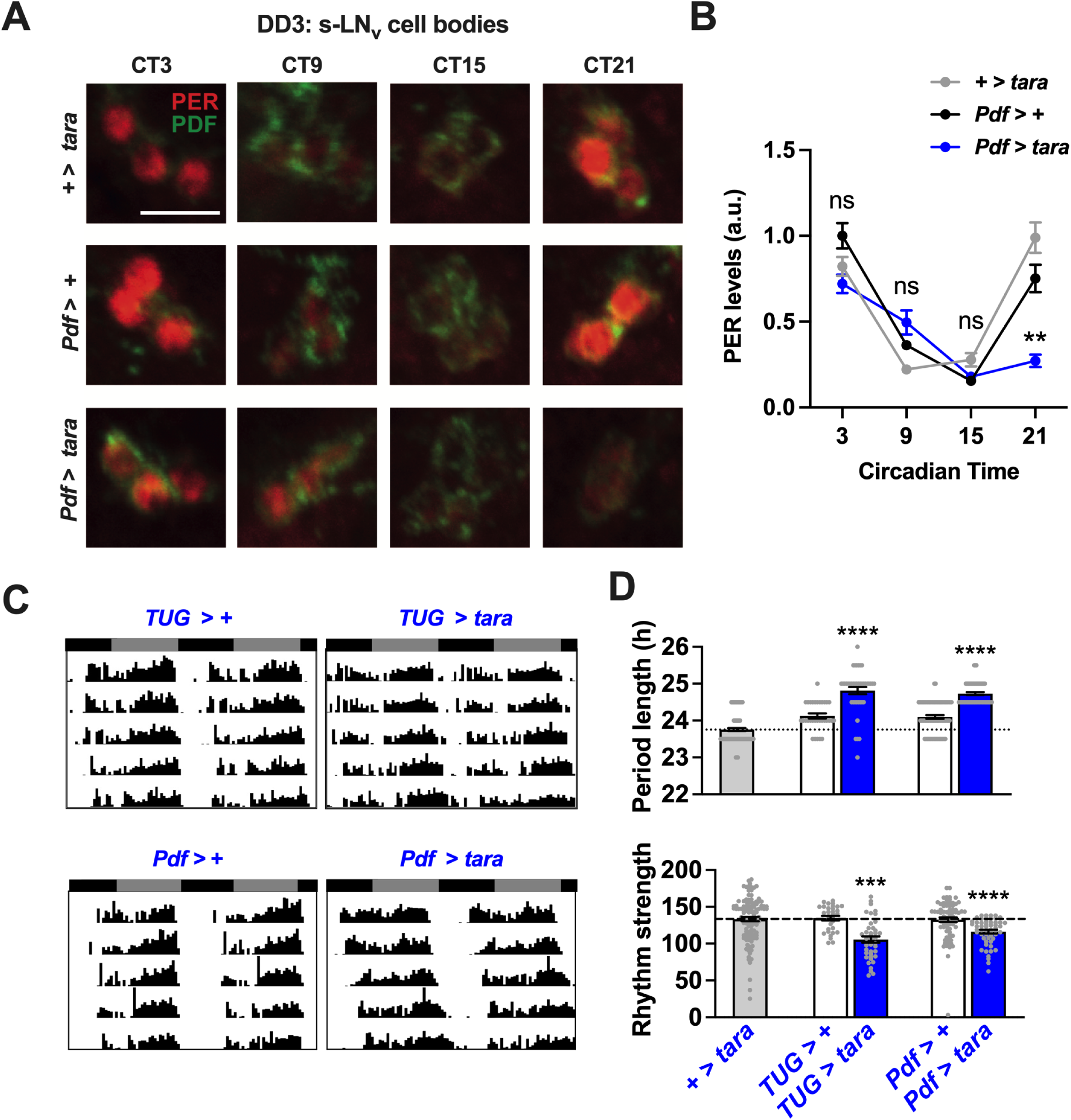
Overexpression of TARA in LN_v_s alters PER cycling and lengthens circadian period. **(A)** Representative expression patterns of PER and PDF in s-LN_v_s of flies overexpressing TARA (*Pdf* > *tara*) and parental controls at the indicated circadian times on the 3^rd^ day in DD. The scale bar is 10 μm. **(B)** PER signal quantification in s-LN_v_s (n = 8-16). **(C)** Representative circadian actograms of male flies overexpressing *tara* in all clock neurons (*TUG tara*) and PDF-positive LN_v_s (*Pdf > tara*) and their parental controls. **(D)** Period length (top) and rhythm strength (bottom) of males of indicated genotypes. The horizontal dotted line represents the mean period length (top) and rhythm strength (bottom) of the common control (+ *tara* RNAi). Mean ± SEM is shown. ****p < 0.0001, ns: not significant; two-way ANOVA followed by Tukey’s post hoc test (B); Kruskal-Wallis test followed by Dunn’s multiple comparisons tests (D).

### Evolutionarily conserved domains in TARA contribute to locomotor rhythmicity

TARA and its mammalian homologs have four conserved domains (Calgaro et al., 2002): CycA binding homology (CAB), SERTA, PHD-Bromo binding (PBB), and C-terminal (CT). To identify the domains essential for circadian regulation, we utilized the CRISPR/Cas9 technique to generate mutants in which the conserved domains were removed from the *tara* genomic locus (Fig. 5A). Deletion of the SERTA domain led to lethality. In contrast, flies in which the CAB, PBB, or CT domain was deleted were viable. We assayed SERTA deletion mutants in trans to the hypomorphic *tara*^s132^ allele and homozygous mutants for all other deletions for circadian locomotor behavior. Compared with *tara*^s132/+^ heterozygous flies, the *tara*^s132/ΔSERTA^ trans-heterozygous mutants exhibited a substantial reduction in rhythm strength, but their circadian period was not significantly altered (Table 3, Fig. 5 B-D). However, given that the s132 allele is hypomorphic, and *tara*^s132/ΔSERTA^ trans-heterozygotes retain some wild-type TARA protein, we cannot rule out the possibility that the SERTA domain plays a role in determining the period length. The lack of period change in *tara*^s132/ΔSERTA^ mutants could be because the more strongly affected mutants were arrhythmic and therefore excluded from the period analysis. Homozygous CAB, PBB, or CT domain deletions had little or no effect on the period length (Table 3, Fig. 5B,C), but they significantly reduced the strength of the locomotor rhythms (Table 3, Fig. 5B,D), suggesting that they also contribute to rhythmic locomotor behavior. Our results indicate that all four conserved domains play a role in maintaining rhythm strength.

**Figure 5.**
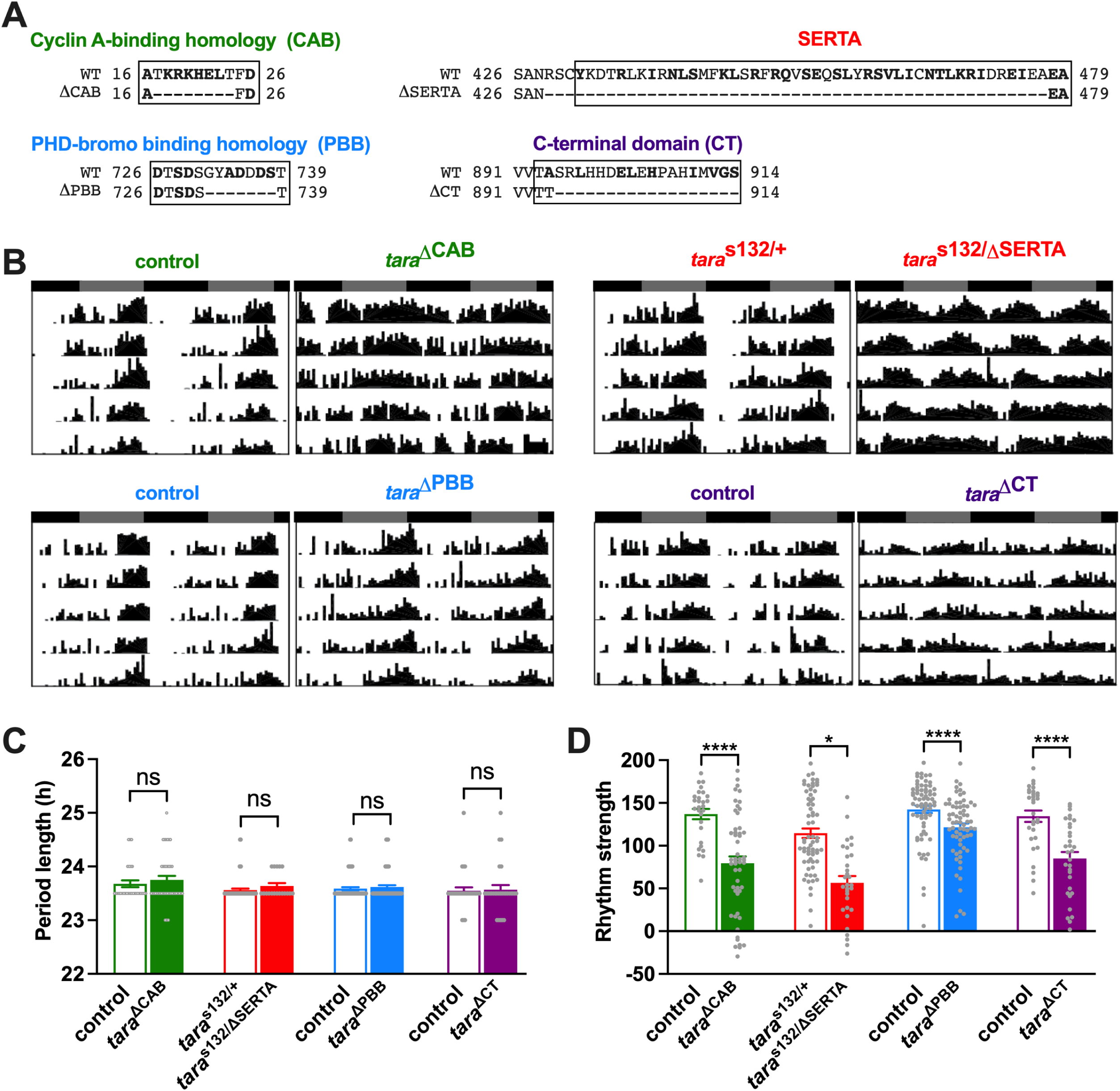
Deletion of conserved TARA domains significantly reduces rhythm strength. **(A)** Amino acid sequences of conserved TARA domains (boxed) and their flanking sequences determined by sequencing the CRISPR mutants. The numbers indicate the positions in the TARA-B sequence. Conserved amino acids are shown in bold. The dashes represent deleted amino acids, and gray indicates substituted amino acids. (**B)** Representative circadian actograms of indicated genotypes. For homozygous domain deletion mutants, we used corresponding homozygous control flies. For *tara*^s132/ΔSERTA^ transheterozygous mutants, *tara*^s132^ mutants were crossed to the control line for *tara*^ΔSERTA^ to generate heterozygous control flies. (**C-D)** Period length (C) and rhythm strength (D) of males of indicated genotypes. N=18-69 for period length, and n=28-70 for rhythm strength. Mean ± SEM is shown. ****p<0.0001, ns: not significant; Kruskal-Wallis test followed by Dunn’s T3 multiple comparisons test (C and D).

**Table 3.** Circadian locomotor rhythm phenotypes of TARA domain deletion mutants.

| **Genotype** | **N** | **% R** | **% WR** | **% AR** | **tau (H) ± Sem** | **Power ± Sem** |
| --- | --- | --- | --- | --- | --- | --- |
| Control | 28 | 100 | 0 | 0 | 23.67 ± 0.06 | 137.0 ± 5.87 |
| *tara*<sup>ΔCAB</sup> | 52 | 71.2 | 5.8 | 23.1 | 23.75 ± 0.07 | 79.4 ± 8.03 |
| *tara*<sup>s132/+</sup> | 66 | 95.5 | 1.5 | 3.0 | 23.56 ± 0.03 | 114.7 ± 5.46 |
| *tara*<sup>s132/ΔSERTA</sup> | 31 | 58.1 | 22.6 | 19.4 | 23.63 ± 0.05 | 56.5 ± 7.76 |
| control | 70 | 97.1 | 1.4 | 1.4 | 23.58 ± 0.03 | 142.5 ± 4.20 |
| *tara*<sup>ΔPBB</sup> | 67 | 94.0 | 1.5 | 4.5 | 23.61 ± 0.03 | 121.5 ± 4.39 |
| control | 30 | 96.8 | 3.2 | 0 | 23.54 ± 0.06 | 134.4 ± 6.59 |
| *tara*<sup>ΔCT</sup> | 31 | 71.6 | 15.6 | 12.5 | 23.56 ± 0.08 | 85.0 ± 7.57 |

N: number of flies; R: rhythmic; WR: weakly rhythmic; AR: arrhythmic.
tau: free-running period; Power: a measure of rhythm strength

### TARA interacts with VRI and PDP1 to modulate *Clk* transcription

Considering the role mammalian homologs of TARA play in transcriptional coregulation (Hsu et al., 2001), our discovery that TARA levels modulate the period length suggests that TARA acts as a transcriptional coregulator of core clock components. We hypothesized that TARA influences VRI activity since *tara* and *vri* mutants exhibit similar circadian phenotypes. Overexpression of *tara* or *vri* in PDF-expressing neurons leads to lengthened period (Table 2, Fig. 4 C,D; Blau and Young, 1999), and both *tara* and *vri* mutants have decreased PDF expression in s-LN_v_s dorsal projections (Fig. 3C,D; Gunawardhana and Hardin, 2017). Consequently, we investigated whether TARA regulates *Clk* transcription repression by VRI. For this purpose, we employed *Drosophila* S2 cells transfected with a *Clk*-*luc* reporter that contains a D-box element, a common binding site for PDP1 and VRI (Cyran et al., 2003). Previous work has shown that VRI binds the same D-box element of the *Clk* promoter as PDP1, acting as a competitive repressor (Cyran et al., 2003). Since repression by VRI would not be evident without *Clk* transcription activation by PDP1, we co-transfected PDP1 and VRI at varying levels of TARA.

A prior study used mammalian HEK293 cells to demonstrate PDP1-dependent *Clk* transcription activation because it was not detectable in S2 cells (Cyran et al., 2003). Intriguingly, we observed PDP1-dependent *Clk* transcription in *Drosophila* S2 cells when TARA was co-transfected with PDP1 (Fig. 6A). Little PDP1 activity was observed without co-transfected TARA, while PDP1 activity increased up to ∼5-fold with rising TARA levels. Transfection of TARA alone did not influence *Clk* transcription (Fig. 6A), in line with its lack of a DNA binding domain and its function as a transcriptional coregulator rather than a transcription factor. VRI co-transfection repressed PDP1-dependent *Clk* transcription, verifying VRI’s role as a competitive repressor of PDP1 (Fig. 6A). The degree of VRI-mediated suppression of *Clk* transcription intensified with increasing levels of TARA. Intermediate TARA levels, which significantly affected PDP1 activity, had little to moderate impact on VRI activity, but higher TARA levels resulted in robust suppression of PDP1 activity by VRI (Fig. 6A). However, since VRI suppressed nearly all PDP1-mediated *Clk* transcription at every level of TARA, the smaller VRI-dependent suppression at lower levels of TARA may be attributed to lower PDP1 activity rather than lower VRI activity.

**Figure 6.**
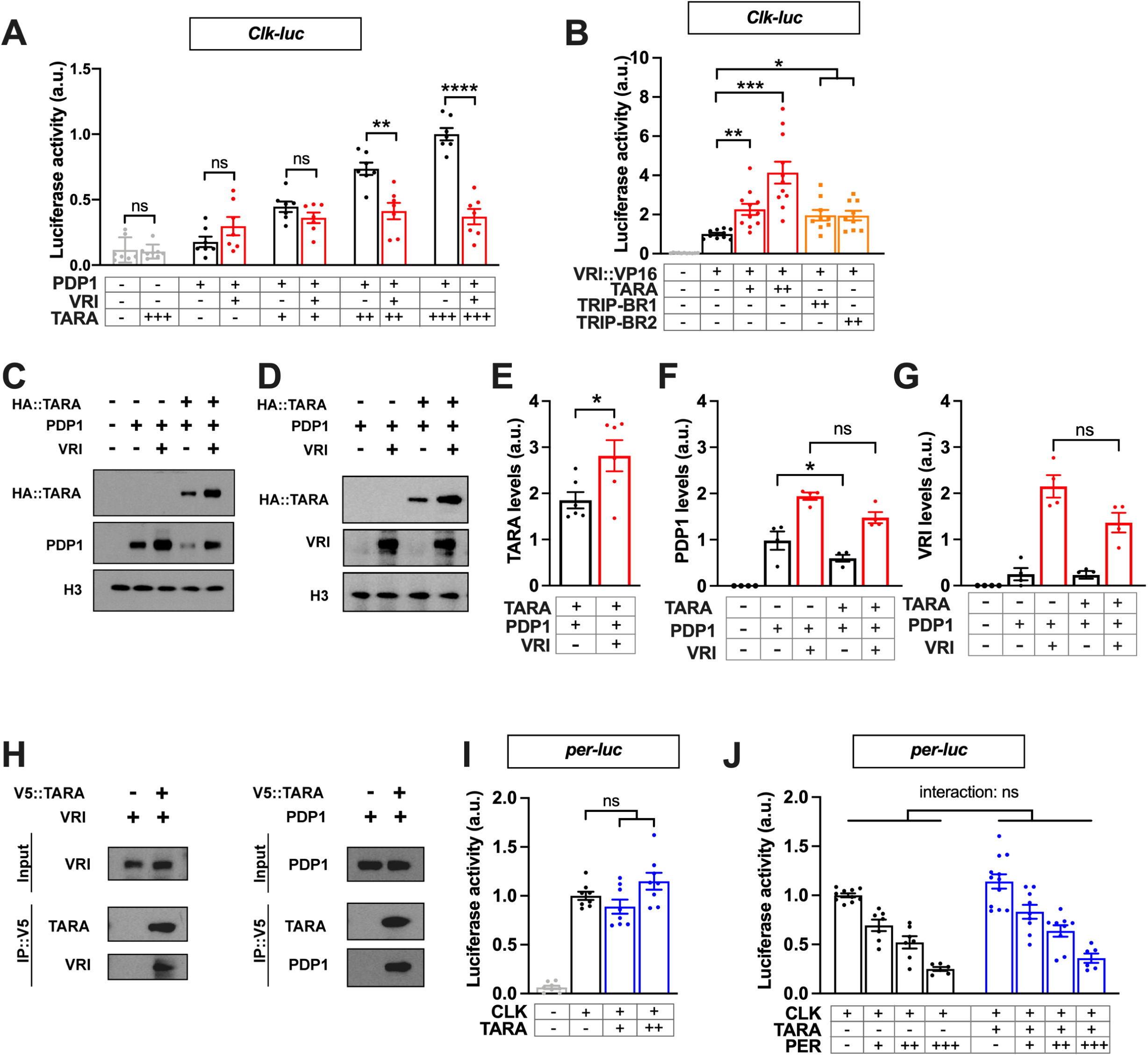
TARA interacts with PDP1 and VRI to regulate *Clk* transcription. **(A)** Luciferase activity of the *Clk*-*luc* reporter in *Drosophila* S2 cells transfected with UAS-*Pdp1* (10 ng), UAS-*vri* (35 ng), and UAS-HA::*tara* (5 ng, 10 ng, or 15 ng) as indicated. Luciferase activity was normalized to the condition that includes UAS-*Pdp1* and 15 ng of UAS-HA::*tara* (+++). **(B)** Luciferase activity of the *Clk*-*luc* reporter in HEK293 cells transfected with CMV-*vri::VP16* (5 ng), CMV-*tara* (10 ng or 30 ng), and CMV-*TRIP-Br1* or CMV-*TRIP-Br2* (30 ng) as indicated. Relative luciferase activity compared to the condition in which CMV-*vri::VP16* alone is transfected is presented. **(C-D).** Representative Western blot of *Drosophila* S2 cells transfected with the *Clk-luc* reporter, UAS-*Pdp1*, UAS-*vri*, and UAS-HA::*tara*, as indicated. Antibodies against HA, PDP1, VRI, and Histone H3 were used. (**E-G)** Quantification of TARA (E), PDP1 (F), and VRI (G) expression levels. **(H)** TARA forms a physical complex with VRI and PDP1 in S2 cells. V5::TARA was immunoprecipitated with an anti-V5 antibody, and antibodies against TARA, PDP1, and VRI were used for Western blotting. **(I)** Luciferase activity of the *per*-*luc* reporter in *Drosophila* S2 cells transfected with UAS-HA::*tara* (10 ng or 30 ng), and UAS-*Clk* (1 ng). Relative luciferase activity compared to the condition in which UAS-*Clk* alone is transfected is presented. **(J)** The assay was conducted as in (I), except with three doses of co-transfected UAS-*per* (2 ng, 5 ng, or 10 ng). The effect of PER levels on *per-luc* activity was significant irrespective of the presence (+) or absence (-) of TARA. None of the pairwise comparisons of the +TARA vs. -TARA conditions at each level of PER revealed a significant difference. Mean ± SEM is shown. ****p<0.0001, ns: not significant; Brown-Forsythe and Welch ANOVA followed by Dunnett’s T3 multiple comparisons test (A, B, and I); Welch’s t-test (E and G); Kruskal-Wallis test followed by Dunn’s multiple comparisons test (F); two-way ANOVA followed by Sidak’s multiple comparisons tests (J).

To investigate the impact of TARA on VRI activity directly without co-transfected PDP1, we utilized VRI fused to the viral VP16 transactivation domain (VRI::VP16), which converts the repressor into a transcriptional activator (Cyran et al., 2003). We were unable to express detectable amounts of VRI::VP16 in *Drosophila* S2 cells, possibly due to lethality caused by its expression. Therefore, we turned to HEK293 cells, as a previous study demonstrated *Clk* transcription by VRI::VP16 using the cell line (Cyran et al., 2003). Consistent with the previous finding, VRI::VP16 promoted *Clk* transcription without co-transfected TARA (Fig. 6B). Importantly, we discovered that co-transfected TARA upregulated VRI::VP16-dependent *Clk* transcription up to ∼4 fold in a dose-dependent manner, demonstrating TARA’s role in enhancing VRI activity. Moreover, TRIP-Br1 and TRIP-Br2, two mammalian homologs of TARA, exhibited a moderate effect on VRI::VP16-dependent *Clk* transcription as well, suggesting a conserved mechanism (Fig. 6B).

To evaluate whether TARA impacts *Clk* transcription by influencing PDP1 and VRI protein levels, we conducted a Western analysis using cell lysates following transfection in S2 cells. Since transcriptional activity occurs in the nucleus, we examined nuclear fractions. Co-transfected TARA did not influence PDP1 or VRI levels (Fig. 6C-G), suggesting that TARA enhances the transcriptional activity of PDP1 and VRI, not their expression or protein stability. Although co-transfected VRI increased TARA and PDP1 levels through an unknown mechanism (Fig. 6C-G), these effects would not account for the observed effects of TARA on *Clk* transcription. Our data support TARA’s function as a transcriptional coregulator of VRI and PDP1.

We then inquired whether the effects of TARA on VRI and PDP1 were mediated by physical interaction. Co-immunoprecipitation (Co-IP) experiments in S2 cells demonstrated that TARA can indeed form a physical complex with VRI or PDP1 (Fig. 6H). Collectively, our results indicate that TARA can physically interact with VRI and PDP1 to enhance the transcriptional suppression of *Clk* by VRI and its activation by PDP1.

### TARA exhibits little or no impact on CLK and PER activity

Our findings indicate that TARA plays a role in the PDP1/VRI feedback loop of the *Drosophila* molecular clock. To ascertain whether TARA also modulates the CLK-CYC/PER-TIM feedback loop, we utilized a *per-luc* reporter in S2 cells and co-transfected CLK with TARA. CLK-mediated transcription of *per* in S2 cells did not require co-transfected TARA, in agreement with previous findings (Darlington et al., 1998), and was unaffected by increasing TARA levels (Fig. 6I).

To investigate whether TARA regulates the suppression of CLK activity by PER, we co-transfected CLK with varying levels of PER in S2 cells, with or without TARA. As previously reported (Darlington et al., 1998), PER repressed CLK-dependent transcription of *per* in a dose-dependent manner (Fig. 6J). Notably, the effect of TARA at each level of PER and the interaction between the effects of TARA and PER were not significant, indicating that PER suppression is independent of TARA (Fig. 6J). Together, our results reveal that TARA exerts minimal impact on CLK’s transcriptional activity and PER’s repressive activity, demonstrating the specificity in TARA’s function as a transcriptional coregulator.

### The SERTA domain is essential for transcriptional coregulation

To investigate the role of TARA’s conserved domains in transcriptional regulation, we generated deletion constructs for each domain: ΔCAB, ΔSERTA, ΔPBB, and ΔCT (Fig. 7A). We transfected full-length or domain-deletion TARA along with *Clk-luc* and PDP1 in S2 cells and conducted Luciferase assays. The results showed that TARA lacking the SERTA domain (ΔSERTA) exerted a markedly diminished effect on PDP1 activity compared to full-length TARA (Fig. 7B), indicating that the SERTA domain is critical for TARA to enhance PDP1’s transcriptional activity. Deletion of the other conserved domains had little effect on PDP1-dependent *Clk* transcription. Next, we examined whether the SERTA domain is essential for TARA’s enhancement of VRI transcriptional activity using VRI::VP16 and HEK293 cells. We discovered that the SERTA domain deletion eliminated TARA’s influence on VRI::VP16 activity (Fig. 7C). To determine the expression levels of deletion constructs and their effects on PDP1 expression, we performed a Western analysis. The results demonstrated that the deletion constructs were expressed at levels similar to or slightly higher than the wild-type TARA (Fig. 7D,E). Furthermore, PDP1 expression was not significantly affected by deletion constructs (Fig. 7D,F), indicating that the reduced *Clk* transcription upon ΔSERTA transfection is not due to diminished PDP1 expression. Taken together, our findings demonstrate that the SERTA domain of TARA is essential for enhancing PDP1 and VRI activity.

**Figure 7.**
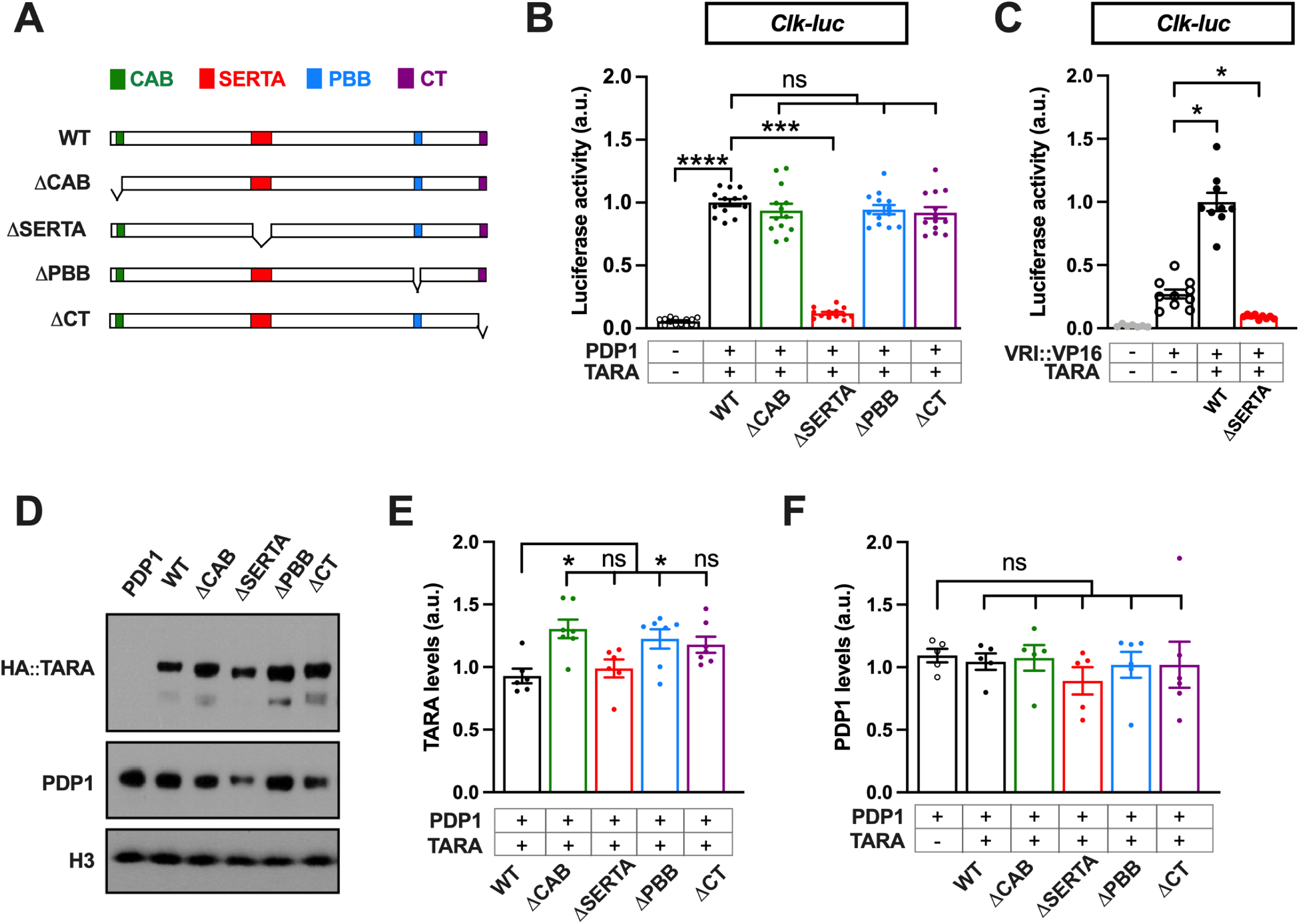
The SERTA domain of TARA is essential for transcriptional coregulation. **(A)** Schematic of TARA domain deletion mutants. **(B)** Luciferase activity of the *Clk*-*luc* reporter in *Drosophila* S2 cells transfected with 30 ng of full-length UAS-HA::*tara* (WT) or HA-tagged TARA domain deletion constructs (ΔCAB, ΔSERTA, ΔPBB, and ΔCT). Luciferase activity compared to the condition in which UAS*-Pdp1* is transfected with full-length TARA is presented. **(C)** Luciferase activity of the *Clk*-*luc* reporter in HEK293 cells transfected with 5 ng of CMV-*vri::VP16* and 30 ng of full-length CMV-*tara* (WT) or a SERTA domain deletion construct (ΔSERTA). Relative luciferase activity compared to the condition in which CMV*-vri::VP16* is transfected with full-length CMV-*tara* is presented. **(D)** Western blot of *Drosophila* S2 cells transfected with a *Clk*-*luc* reporter with 30 ng of full-length UAS-HA::*tara* (WT) or HA-tagged domain deletion constructs (ΔCAB, ΔSERTA, ΔPBB, and ΔCT) and 15 ng UAS-*Pdp1* as indicated. (**E-F**) Quantification of TARA (E) and PDP1 (F) expression levels. Mean ± SEM is shown. Mean ± SEM is shown. ****p < 0.0001, ns: not significant; Kruskal-Wallis followed by Dunn’s multiple comparisons test relative to a control condition (B,C,E, and F).

### *tara* genetically interacts with *vri* and *Pdp1* to modulate the molecular clock *in vivo*

Since we discovered that TARA enhanced the transcriptional activity of VRI and PDP1 in cultured cells, we examined whether *tara* interacts with *vri* and *Pdp1* to modulate the molecular clock *in vivo*. We overexpressed *vri* or *Pdp1* in PDF+ clock neurons, with or without *tara* co-overexpression. We observed a pronounced increase in period length when *vri* was overexpressed in LN_v_s (Table 4, Fig. 8), consistent with previous findings (Blau and Young, 1999). Although overexpressing *tara* alone also significantly increased period length, co-overexpressing both *tara* and *vri* did not further increase the period compared to overexpressing *vri* alone (Table 4, Fig. 8), indicating a genetic interaction between *tara* and *vri*. TARA may lengthen the circadian period by enhancing VRI activity when VRI is present at normal levels. However, when VRI levels are elevated, additional TARA may not further amplify VRI’s period-lengthening effect.

**Figure 8.**
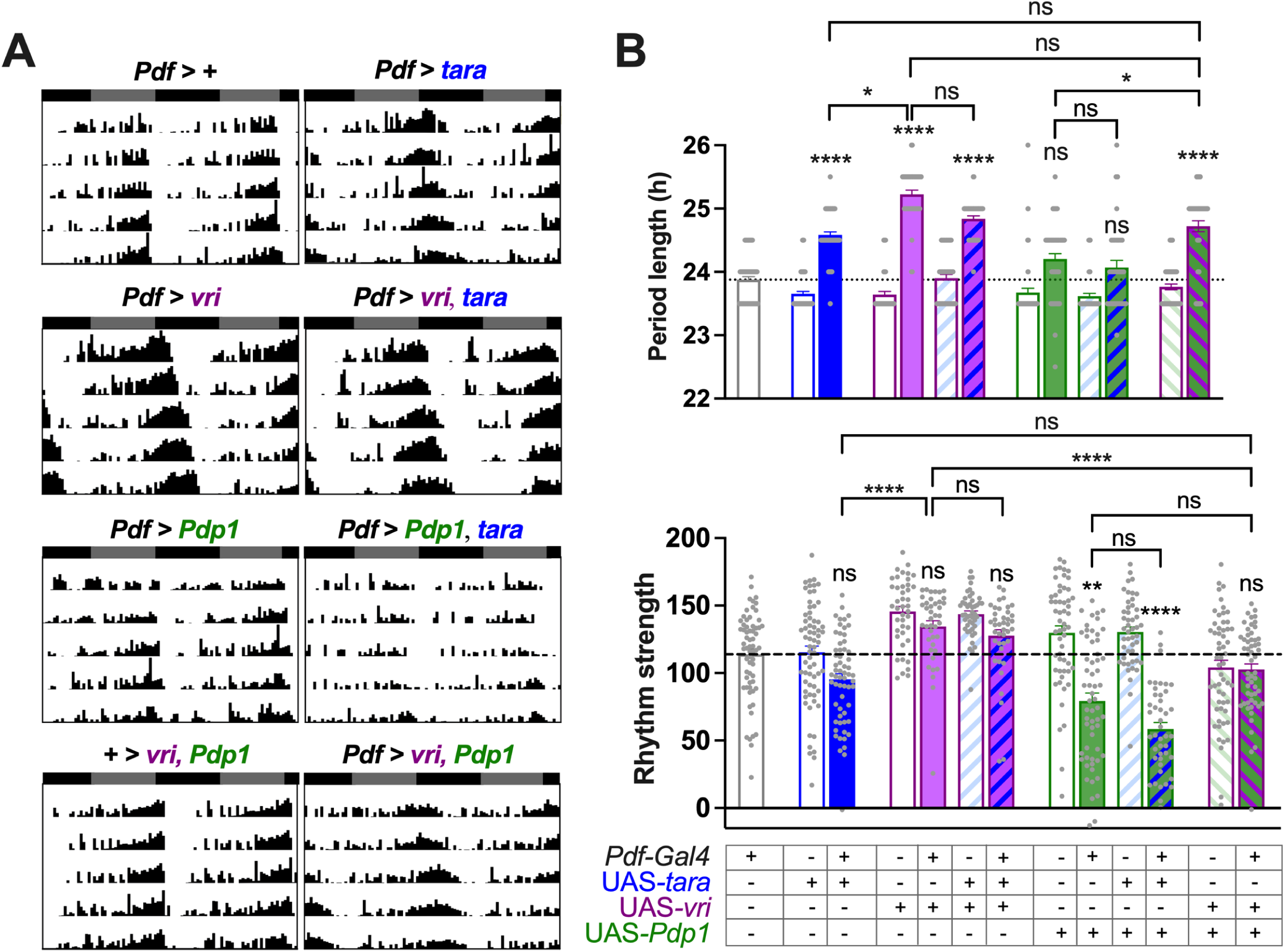
*tara* regulates the clock speed through genetic interactions with *vri* and *Pdp1*. (**A**) Representative circadian actograms of males of indicated genotypes. (**B**) Period length (top) and rhythm strength (bottom) of indicated genotypes. N = 36-64 for period length, n = 40-67 for rhythm strength. The horizontal dotted line represents the mean period length (top) and rhythm strength (bottom) for the common parental control (*Pdf* > +). Mean ± SEM is shown. ****p < 0.0001, ns: not significant; Kruskal-Wallis followed by Dunn’s multiple comparisons test.

**Table 4.** Circadian locomotor rhythm phenotypes of TARA, VRI, and PDP1 overexpression in PDF+ neurons.

| **Genotype** | **N** | **% R** | **% WR** | **% AR** | **tau (h) ± SEM** | **Power ± SEM** |
| --- | --- | --- | --- | --- | --- | --- |
| *Pdf*> + | 64 | 95.2 | 3.2 | 1.6 | 23.88 ± 0.06 | 113.8 ± 3.94 |
| + > *tara* | 67 | 92.5 | 6 | 1.5 | 23.61 ± 0.03 | 115.3 ± 4.59 |
| *Pdf*> *tara* | 61 | 93.5 | 4.9 | 1.6 | 24.57 ± 0.05 | 95.4 ± 4.15 |
| + > *vri* | 42 | 100 | 0 | 0 | 23.64 ± 0.05 | 145.6 ± 3.85 |
| *Pdf*> *vri* | 40 | 97.5 | 2.5 | 0 | 25.23 ± 0.07 | 134.5 ± 4.21 |
| + > *vri, tara* | 46 | 100 | 0 | 100 | 23.90 ± 0.06 | 143.6 ± 2.39 |
| *Pdf*> *vri, tara* | 40 | 95 | 5 | 0 | 24.84 ± 0.05 | 127.6 ± 4.50 |
| + > *Pdp1* | 55 | 96.4 | 1.8 | 1.8 | 23.68 ± 0.07 | 129.8 ± 5.22 |
| *Pdf*> *Pdp1* | 59 | 67.8 | 18.6 | 13.6 | 24.21 ± 0.08 | 79.4 ± 5.81 |
| + > *Pdp1, tara* | 47 | 97.9 | 2.1 | 0 | 23.62 ± 0.04 | 130.4 ± 3.79 |
| *Pdf*> *Pdp1, tara* | 44 | 59.1 | 22.7 | 18.2 | 24.07 ± 0.11 | 58.5 ± 4.80 |
| + > *Pdp1, vri* | 56 | 93 | 3.5 | 3.5 | 23.76 ± 0.04 | 104.3 ± 5.27 |
| *Pdf*> *Pdp1, vri* | 55 | 94.6 | 3.6 | 1.8 | 24.72 ± 0.09 | 102.7 ± 4.16 |

N: number of flies; R: rhythmic; WR: weakly rhythmic; AR: arrhythmic.
tau: free-running period; Power: a measure of rhythm strength.

Overexpression of *Pdp1* did not lead to a significant change in the period length (Table 4, Fig. 8), consistent with a previous finding (Zheng et al., 2009). Although overexpressing *tara* in PDF+ neurons led to a significantly longer period, co-overexpression with *Pdp1* eliminated TARA’s impact on the period length (Table 4, Fig. 8), demonstrating that *tara* and *Pdp1* genetically interact to regulate the pace of the molecular clock.

Overexpression of *tara*, *vri,* alone or in combination with *tara* had minor effects on the robustness of circadian rhythms (Table 4, Fig. 8). In contrast, overexpression of *Pdp1* significantly weakened the strength of locomotor rhythms, consistent with previous findings (Benito et al., 2007; Zheng et al., 2009). Co-overexpression of *tara* and *Pdp1* further decreased the rhythm strength, but the decrease was not statistically significant (Table 4, Fig. 8).

The effect of *tara* overexpression on the circadian period may appear more similar to that of *vri* overexpression than *Pdp1* overexpression, as both *tara* and *vri* overexpression markedly lengthen the period while *Pdp1* overexpression does not have a significant effect. However, it is important to note that this observation does not necessarily indicate that TARA has a greater impact on the transcriptional activity of VRI compared to PDP1. Considering that TARA enhanced VRI and PDP1 activity in S2 cells, the impact of *tara* overexpression on locomotor rhythms *in vivo* may reflect the combined effects of enhancing both VRI and PDP1 activity.

Indeed, when we co-overexpressed *vri* and *Pdp1* in PDF+ clock neurons, we observed an increase in period length similar to that of *tara* overexpression (Table 4, Fig. 8). Co-overexpressing *vri* and *Pdp1* also resulted in a rhythm strength comparable to that of overexpressing *tara*. Collectively, our findings indicate that *tara* interacts with *vri* and *Pdp1* to modulate the circadian period and rhythm strength in *Drosophila*.

## Discussion

The present study identifies TARA as a novel regulatory component of the interlocked transcription-translation feedback loops in the *Drosophila* molecular clock. Whereas the primary feedback loop involving CLK-CYC and PER-TIM in the *Drosophila* circadian clock has been studied extensively (Allada and Chung, 2010; Hardin, 2011; Tataroglu and Emery, 2015; Patke et al., 2020), the secondary loop, including VRI and PDP1, is relatively understudied (Blau and Young, 1999; Cyran et al., 2003; Glossop et al., 2003; Benito et al., 2007; Zheng et al., 2009; Ling et al., 2012; Gunawardhana and Hardin, 2017; Gunawardhana et al., 2021). Our transcriptional assays reveal that TARA enhances the activity of both the negative and positive arms of the secondary feedback loop. TARA has minimal effects on PER and CLK activity, suggesting that TARA plays a more critical role in the secondary loop than the primary loop. Consistent with the transcriptional assay results, our behavioral data demonstrate that *tara* genetically interacts with *vri* and *Pdp1* to regulate the pace of the *Drosophila* circadian clock.

*tara* mutants display markedly impaired locomotor rhythmicity despite robust PER cycling in s-LNvs, suggesting that TARA plays an additional role downstream of the s-LNv molecular clock. TARA may regulate the output of the s-LN_v_ or other clock neurons. Future studies examining PER cycling in other clock neuron clusters may reveal whether communications between s-LN_v_s and other clock neurons or those between clock and non-clock neurons are disrupted. Our findings are reminiscent of previous results showing that VRI and PDP1 play critical roles in the molecular clock as well as downstream of the s-LNv clock (Blau and Young, 1999; Cyran et al., 2003; Glossop et al., 2003; Benito et al., 2007; Zheng et al., 2009; Ling et al., 2012; Gunawardhana and Hardin, 2017; Gunawardhana et al., 2021), supporting the view that TARA functions with VRI and PDP1 to regulate circadian rhythms.

Like VRI and PDP1, TARA has developmental roles (Calgaro et al., 2002; Manansala et al., 2013). Although PDF expression patterns in the s-LN_v_ dorsal projections in *tara* mutants do not suggest gross morphological defects, we cannot rule out subtle developmental defects or reduced adult plasticity in the complexity of the projections (Fernandez et al., 2008). However, even dramatic changes in the s-LN_v_ morphology and a loss of adult plasticity do not impair rhythmic behavior under standard constant-temperature conditions (Fernandez et al., 2020). Thus, it is unlikely that the reduced rhythmicity of *tara* mutants results from s-LN_v_ morphological defects. We previously reported that TARA promotes sleep through its interaction with CycA and Cdk1 in a subset of neurons in the *pars lateralis*, a region considered analogous to the mammalian hypothalamus (Afonso et al., 2015b). Collectively, our findings suggest that TARA regulates circadian rhythms and sleep by interacting with different molecular partners in distinct neuronal populations.

Given that TARA enhances both the activator and suppressor of *Clk* transcription, *tara* overexpression might be expected to have little or no effect on the circadian period. However, *vri* overexpression leads to a markedly long period, while *Pdp1*overexpression does not significantly affect the period. Therefore, the combined effect of enhancing both VRI and PDP1 is expected to be a longer period compared to control flies. Indeed, we find that *tara* overexpression lengthens the circadian period, mirroring the consequence of simultaneously overexpressing *vri* and *Pdp1*. It is unclear why *vri* overexpression affects the clock speed more than *Pdp1* overexpression. A previous study demonstrated that VRI expression peaks a few hours before PDP1 through an as-yet unidentified mechanism (Cyran et al., 2003). The earlier accumulation of VRI may be linked to a more potent effect of *vri* overexpression on the circadian period.

The components of the primary feedback loop in the molecular clock are highly conserved between flies and mammals (Allada and Chung, 2010; Hardin, 2011; Tataroglu and Emery, 2015; Patke et al., 2020). Mammalian CLK, BMAL1 (a CYC homolog), and PER1/2/3 perform similar functions as their *Drosophila* orthologs, while CRYPTOCHROME (CRY)1/2 are functionally equivalent to *Drosophila* TIM. However, the functions of the secondary feedback loop involving VRI and PDP1 in *Drosophila* are performed by more complex, multi-layered feedback loops in mammals. The layered loops include nuclear receptors, REV-ERBα/β (also known as NR1D1/2), and Retinoid-related Orphan Receptor (ROR)α/β/δ, which competitively bind the RRE element in the promotor of RRE-containing genes, including *BMAL1*, to activate and suppress its transcription (Preitner et al., 2002; Ueda et al., 2002; Sato et al., 2004). REV-ERBs and RORs are not homologs of VRI and PDP1, but mammalian homologs of VRI and PDP1, named Nuclear Factor Interleukin-3-Regulated Protein (NFIL3, also known as E4BP4) and D-BOX BINDING PROTEIN (DBP), respectively, also participate in the interlocked feedback loops. Like their *Drosophila* homologs, NFIL3 and DBP competitively bind the D-box element to inhibit and activate the transcription of D-box-containing genes, including RORs (Mitsui et al., 2001). RORs activate *NFIL3* transcription, while NFIL3 inhibits the transcription of *ROR*s, forming a feedback loop (Mitsui et al., 2001; Gachon et al., 2004; Ueda et al., 2005). Taken together, REV-ERBs and NFIL3 suppress *BMAL1* transcription, while RORs and DBP activate *BMAL1* transcription. These findings suggest that VRI performs the combined functions of REV-ERBs and NFIL3, while PDP1 performs the combined functions of RORs and DBP. Initial studies demonstrating that VRI and PDP1 regulate *Clk* transcription were performed in mammalian HEK293 cells because PDP1 did not activate *Clk* transcription in S2 cells (Cyran et al., 2003). Our data show that TARA is the missing factor necessary for *Clk* transcription by PDP1 in *Drosophila* S2 cells. HEK293 cells may express sufficient levels of TRIP-Br proteins, the mammalian homologs of TARA, to allow PDP1 activity at the *Clk* promoter. These considerations and our finding that TRIP-Br1/2 enhance VRI activity in HEK293 cells raise the intriguing possibility that TRIP-Br proteins regulate the mammalian circadian period through their interactions with NFIL3 and DBP for the transcription of D-box-containing genes.

TARA and TRIP-Br proteins contain four conserved domains (Calgaro et al., 2002; Manansala et al., 2013). The function of the SERTA domain is not well understood. The present study uncovers a novel role of the SERTA domain in enhancing PDP1 and VRI activity. We also find that TARA is a critical regulator of rhythm amplitude and that all four domains contribute to robust locomotor rhythms. The reduced rhythmicity in the deletion mutants is unlikely to be due to reduced stability of TARA with domain deletions since deletion constructs were expressed at levels comparable to or slightly higher than wild-type TARA in S2 cells (Fig. 7D,E). It would be interesting to investigate further the molecular functions of the conserved domains and their role in circadian rhythms.

TARA is ubiquitously expressed in neurons (Afonso et al., 2015b), but what regulates TARA expression or activity is largely unknown. Notably, chromatin immunoprecipitation tiling array assays (Abruzzi et al., 2011) detected cyclic CLK binding to a promoter region of *tara*, suggesting that it is a direct transcriptional target of CLK. Although we did not observe daily cycling of TARA protein levels in s-LN_v_s (Afonso et al., 2015a), TARA activity may exhibit circadian cycling, since its partners VRI and PDP1 undergo circadian cycling. TARA levels or activity may also respond to changing environmental and internal conditions, such as the metabolic state. Earlier findings suggest that mammalian homologs of TARA are involved in metabolic regulation. TRIP-Br2 is involved in lipid and oxidative metabolism in adult mice (Liew et al., 2013), and prolonged nutrient starvation dramatically decreases TRIP-Br3 to induce apoptosis (Li et al., 2015). Circadian regulation is closely linked to metabolism (Bailey et al., 2014; Asher and Sassone-Corsi, 2015; Guan and Lazar, 2021), and TARA may play a role in integrating circadian and metabolic regulation. Further studies will need to be performed to examine the potential role of TARA in linking circadian rhythms and metabolism.

Our findings reveal a novel role for *tara* in regulating the pace of the circadian clock through its interaction with *vri* and *Pdp1* in the secondary feedback loop of the *Drosophila* circadian clock. They provide a rationale for investigating the role of the TRIP-Br proteins in regulating circadian rhythms and their potential interaction with NFIL3 and DBP in the mammalian molecular clock.

## Acknowledgments

This work was supported by grants from the National Institutes of Health (R01NS086887 and R01NS109151 to K.K.) and funds from Thomas Jefferson University Synaptic Biology Center (to K.K.). We thank the Bloomington *Drosophila* Stock Center and National Institute of Genetics for fly stocks; *Drosophila* Genomics Research Center (supported by NIH grant 2P40OD010949-10A1) for vectors; Drs. Amita Sehgal, Paul Hardin, Justin Blau, and Jit Kong Cheong for antibodies and constructs; M. Boudinot and Dr. Francois Rouyer for the Faas software; Luke Kim and William Wei for technical assistance; and Drs. Yohei Kirino, Joanna Chiu, Aaron Haeusler, Eun Young Kim, Sang Hyuk Lee, and James Jaynes, and members of the Koh lab for helpful discussions.

